# Development of new cowpea (*Vigna unguiculata*) mutant genotypes, analysis of their agromorphological variation, genetic diversity and Population structure

**DOI:** 10.1101/2020.05.22.109041

**Authors:** Made Diouf, Sara Diallo, François Abaye Badiane, Oumar Diack, Diaga Diouf

**Affiliations:** Laboratoire Campus de Biotechnologies Végétales, Département de Biologie Végétale, Faculté des Sciences et Techniques, Université Cheikh Anta Diop, Dakar-Fann, Code Postal 10700, Dakar, Sénégal; Faculté des Sciences et Technologies de l’Education et de la Formation, Université Cheikh Anta Diop, Dakar-Fann, Code Postal 10700, Dakar, Sénégal

**Keywords:** Cowpea, *Vigna unguiculata*, Gamma rays, Induced mutagenesis, Plant breeding, Agromorphological characterization, ISSR

## Abstract

Cowpea is one of the most important legume grain in the SubSaharian region of Africa used for human consumption and animal feeding but its production is hampered by biotic and abiotic constraints raising the need to broaden its genetic basis. For this purpose, the seeds of two cowpea varieties Melakh and Yacine were respectively irradiated with 300 and 340 Gy. The developed mutant populations were agromorphologically characterized from M5 to M7 while the genetic diversity of the last were evaluated using 13 ISSR markers. Based on agromorphological characterization, variation of flower color, pod length, seed coat color and seed weight with respectively 78.01, 68.29, 94.48 and 57.58% heritability were recorded in the mutant lines. PCA analyses allowed to identify the elite mutants based on their agromorphological traits while Pearson’s correlation results revealed a positive correlation between yield component traits. Three subpopulations were identified through STRUCTURE analyses but assignment of the individuals in each group was improved using DAPC. Analysis of Molecular Variance revealed that the majority (85%) of the variance rather existed within group than among (15%) group. Finally, our study allowed to select new promising mutant genotypes which could be tested for multi local trials to evaluate their agronomic performance.

## Introduction

Cowpea [*Vigna unguiculata* (L.) Walp. 2n = 2x =22] is an important crop legume for tropical and subtropical regions grown in Africa, Southern Europe, Latin America, Southeast Asia, and southwestern regions of North America on 12 496 305 hectares (http://www.fao.org/faostat/en/#data/QC/visualize). Its production is estimated to 7 233 408 tons, Nigeria, Niger, Burkina Faso, Ghana, United Republic of Tanzania, Myanmar, Mali, Cameroon, Sudan (including Sudan and South Soudan) and Kenya, are the top producers in the world. In the Sahelian part of Africa, the crop plays a major role in human nutrition. For instance, the fresh seeds, grilled on wood fire, are consumed in Senegal while the dry seeds are used in a wide range of meal compositions. The young leaves cooked as spinach are eaten in Eastern and Southern Africa while the hay as well as the seed are used for livestock feeding in several African countries (for review see [1]). Compared to others legumes, its seed contains higher amount of proteins which is estimated to 25% and a substantial amount of minerals and vitamins based on the recent work published by [2], raising cowpea as a valuable crop to fight malnutrition for the low-income farmers. Based on the estimation realized by [3], cowpea which establishes a symbiosis with *Bradyrhizobium*, is a good nitrogen fixing crop (70 to 350 kg nitrogen per hectare) leading to soil fertility improvement. Despite its importance, the production of the crop is hampered by a wide range of biotic (virus, bacteria, fungi, parasitic weeds and nematodes) and abiotic (drought, heat, etc.) constraints [1, 4]. This susceptibility to a wide range of biotic and abiotic stresses is attributed to the narrow genetic basis of the crop due to single domestication event and its self-pollinating pattern of reproduction [5]. To overcome these constraints, the genetic diversity hidden into the existing germplasms which contains relevant agronomic traits have been exploited during the past decades to increase the production of crops through the development of outstanding cultivars, based on methods such as pure line selection, mass selection, pedigree breeding, single seed descent and back crossing [6]. Despite these efforts, the genetic basis of the realized lines is still narrow which arise the need to develop novel outstanding varieties particularly in the era of climate change. For this purpose, technique such as mutagenesis is a valuable tool to induce genetic variation for cultivar improvements. Mutagenesis has widely been used for the past seventy years in order to improve many economically important crops leading to the selection of 3,320 varieties which are superior to the natural cultivar in term of productivity, resistance to biotic and/or abiotic constraints (https://mvd.iaea.org/#!Search). According to this database, only 16 cowpea varieties were bred using gamma ray mutation techniques and 5 of them were selected from Africa precisely in Kenya, Zambia and Zimbabwe.

Gamma rays are ionizing radiation which deeply penetrate into the cells of the target tissues where they interact with molecules to generate reactive oxygen species (ROS) causing base substitutions, genome rearrangements such as insertions, deletions, inversions and translocations [7]. The base substitution caused by ROS is due to the conversion of guanines into 8-oxo-Gs which induces mispairings with adenine while genome rearrangements are caused by error-prone non homologous end joining (NHEJ) rather than error-free homologous recombination which result from double-strand breaks (DSBs). When DSBs and NHEJ occur in several genomic regions, they create favorable conditions for copy number variations (CNVs), presence/absence variations (PAVs) and translocations [8, 7]. Presently, it is well documented that these genetic modifications induce agro morphological variations affecting plant height, growth habit, number of leaves per plant, leaves color, number of branches per plant, days to first flower, flower color, flowering ability, maturity, number of pods per plant, number of seed per plant, pod and seed coat color, seed eye color, weight of 100 seeds and tolerance to *Maruca vitrata* pod borer which were observed among cowpea mutants [9,10,11] [12,13,14]. In view of these variations, several research teams attempted to characterize cowpea mutant population using morphological traits, yield components [15,16,10,12,13,17] and recently seed storage proteins [10, 12]. To overcome the limits of using these traits, Random amplified polymorphic DNA (RAPD; [10]) and inter-simple sequence repeat (ISSR; [10, 12]) were recently used to understand the genetic organization of some cowpea mutant populations. of these markers. The analysis of ISSR has generated more informative results, since these sequences are abundant, widely distributed across the eukaryotic genome, highly reproducible and use SSR as primers allowing the amplification of inter SSR region [18]. ISSR are useful to study genetic diversity, genome mapping or evolutionary biology in many crop species and they overcome the limitations of other markers, such as low reproducibility of RAPD and high cost of AFLP (Amplified fragment length polymorphic) and the use a single primer during the amplification processes [18, 19]. ISSR combine the advantages of SSR, AFLP and RAPD markers which do not require prior genome sequence information and are efficient to detect genetic variation among cowpea varieties [20, 21] or mutant lines [10, 12]. ISSR primers can be unanchored with 1 to 4 degenerate nucleotides at 3’ or 5’ end to avoid their slippery within the repeat units and smears apparition during amplification but previous studies showed that primers anchored at 3’ gave more clear bands [22, 12].

The aim of this study was to broaden the genetic basis of cowpea using gamma irradiation technique specifically to develop mutant populations for which their genetic diversity and allelic richness were assessed and to use the generated information to select new elite genotypes.

## Materials and Methods

### Plant materials and gamma irradiation

Two inbred cowpea varieties Melakh and Yacine (Table 1) widely cultivated in Senegal were selected from the national germplasm and used in this study [23]. They belong to the early maturity group which reach physiological maturity 64 days after sowing (DAS) under well-watered conditions [24, 25]. In total 216 dry and healthy seeds for each variety Melakh and Yacine were exposed respectively to 300 and 340 Gy of gamma ray. The irradiation was performed at the International Atomic Energy Agency (IAEA), Agriculture and Biotechnology Laboratory, A-2444 Seibersdorf, Austria using a cobalt 60 source Gammacell Model No. 220. The control seeds were not exposed to gamma irradiation.

**Table 1:**
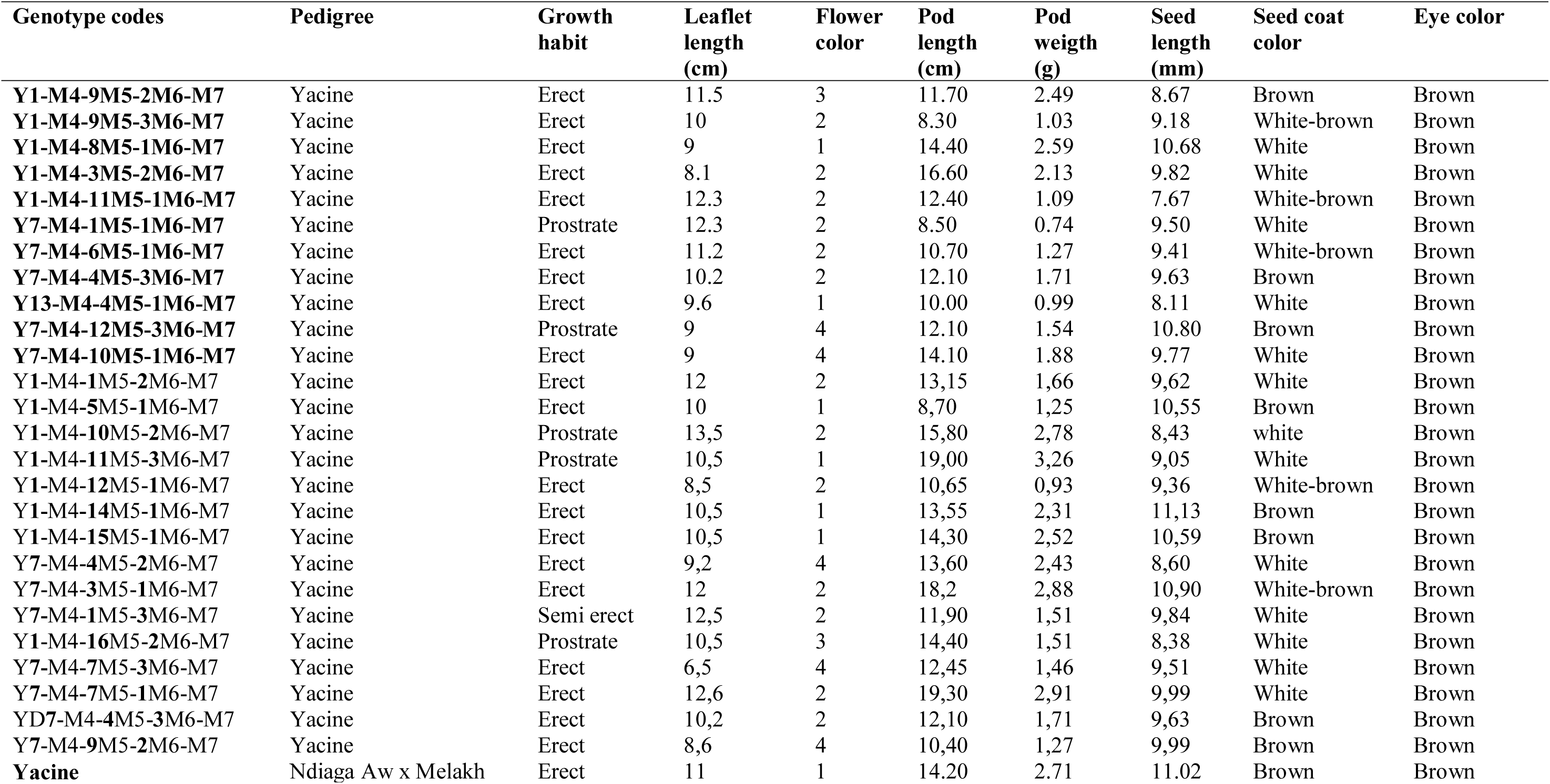
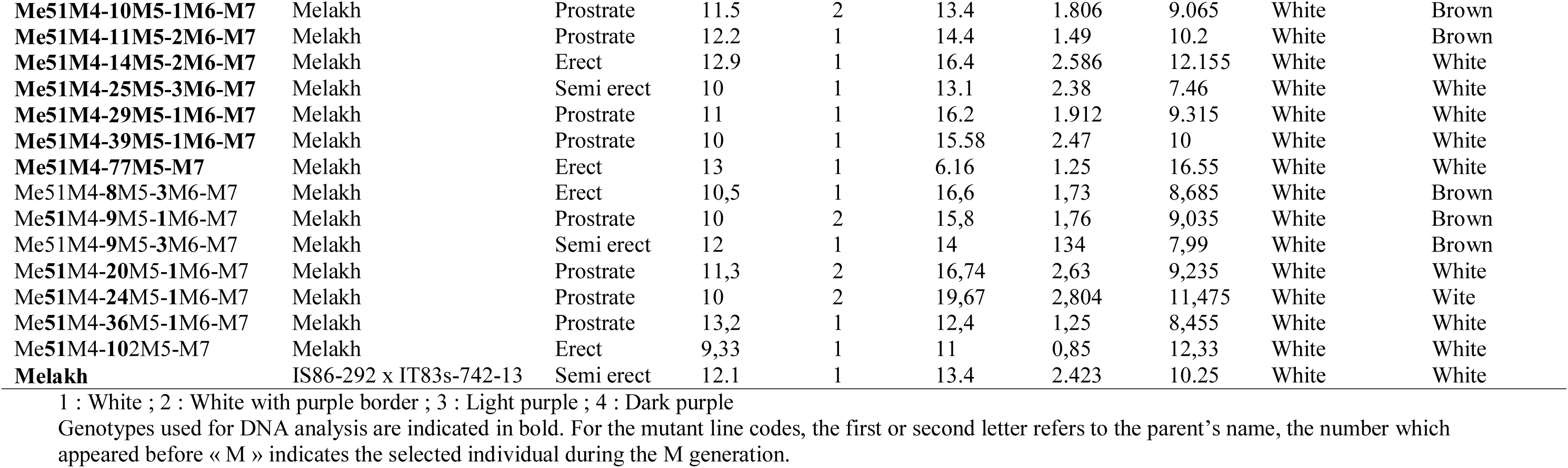
Agromorphological characters and pedigree of 40 cowpea mutants and their parent used in this study.

| Genotype codes | Pedigree | Growth habit | Leaflet length (cm) | Flower color | Pod length (cm) | Pod weight (g) | Seed length (mm) | Seed coat color | Eye color |
| --- | --- | --- | --- | --- | --- | --- | --- | --- | --- |
| **Y1-M4-9M5-2M6-M7** | Yacine | Erect | 11.5 | 3 | 11.70 | 2.49 | 8.67 | Brown | Brown |
| **Y1-M4-9M5-3M6-M7** | Yacine | Erect | 10 | 2 | 8.30 | 1.03 | 9.18 | White-brown | Brown |
| **Y1-M4-8M5-1M6-M7** | Yacine | Erect | 9 | 1 | 14.40 | 2.59 | 10.68 | White | Brown |
| **Y1-M4-3M5-2M6-M7** | Yacine | Erect | 8.1 | 2 | 16.60 | 2.13 | 9.82 | White | Brown |
| **Y1-M4-11M5-1M6-M7** | Yacine | Erect | 12.3 | 2 | 12.40 | 1.09 | 7.67 | White-brown | Brown |
| **Y7-M4-1M5-1M6-M7** | Yacine | Prostrate | 12.3 | 2 | 8.50 | 0.74 | 9.50 | White | Brown |
| **Y7-M4-6M5-1M6-M7** | Yacine | Erect | 11.2 | 2 | 10.70 | 1.27 | 9.41 | White-brown | Brown |
| **Y7-M4-4M5-3M6-M7** | Yacine | Erect | 10.2 | 2 | 12.10 | 1.71 | 9.63 | Brown | Brown |
| **Y13-M4-4M5-1M6-M7** | Yacine | Erect | 9.6 | 1 | 10.00 | 0.99 | 8.11 | White | Brown |
| **Y7-M4-12M5-3M6-M7** | Yacine | Prostrate | 9 | 4 | 12.10 | 1.54 | 10.80 | Brown | Brown |
| **Y7-M4-10M5-1M6-M7** | Yacine | Erect | 9 | 4 | 14.10 | 1.88 | 9.77 | White | Brown |
| Y1-M4-1M5-2M6-M7 | Yacine | Erect | 12 | 2 | 13,15 | 1,66 | 9,62 | White | Brown |
| Y1-M4-5M5-1M6-M7 | Yacine | Erect | 10 | 1 | 8,70 | 1,25 | 10,55 | Brown | Brown |
| Y1-M4-10M5-2M6-M7 | Yacine | Prostrate | 13,5 | 2 | 15,80 | 2,78 | 8,43 | white | Brown |
| Y1-M4-11M5-3M6-M7 | Yacine | Prostrate | 10,5 | 1 | 19,00 | 3,26 | 9,05 | White | Brown |
| Y1-M4-12M5-1M6-M7 | Yacine | Erect | 8,5 | 2 | 10,65 | 0,93 | 9,36 | White-brown | Brown |
| Y1-M4-14M5-1M6-M7 | Yacine | Erect | 10,5 | 1 | 13,55 | 2,31 | 11,13 | Brown | Brown |
| Y1-M4-15M5-1M6-M7 | Yacine | Erect | 10,5 | 1 | 14,30 | 2,52 | 10,59 | Brown | Brown |
| Y7-M4-4M5-2M6-M7 | Yacine | Erect | 9,2 | 4 | 13,60 | 2,43 | 8,60 | White | Brown |
| Y7-M4-3M5-1M6-M7 | Yacine | Erect | 12 | 2 | 18,2 | 2,88 | 10,90 | White-brown | Brown |
| Y7-M4-1M5-3M6-M7 | Yacine | Semi erect | 12,5 | 2 | 11,90 | 1,51 | 9,84 | White | Brown |
| Y1-M4-16M5-2M6-M7 | Yacine | Prostrate | 10,5 | 3 | 14,40 | 1,51 | 8,38 | White | Brown |
| Y7-M4-7M5-3M6-M7 | Yacine | Erect | 6,5 | 4 | 12,45 | 1,46 | 9,51 | White | Brown |
| Y7-M4-7M5-1M6-M7 | Yacine | Erect | 12,6 | 2 | 19,30 | 2,91 | 9,99 | White | Brown |
| YD7-M4-4M5-3M6-M7 | Yacine | Erect | 10,2 | 2 | 12,10 | 1,71 | 9,63 | Brown | Brown |
| Y7-M4-9M5-2M6-M7 | Yacine | Erect | 8,6 | 4 | 10,40 | 1,27 | 9,99 | Brown | Brown |
| **Yacine** | Ndiaga Aw x Melakh | Erect | 11 | 1 | 14.20 | 2.71 | 11.02 | Brown | Brown |

**Table 1: Agro-morphological characters and pedigree of 40 cowpea mutants and their parent used in this study (continued).**
| Genotype codes | Pedigree | Growth habit | Leaflet length (cm) | Flower color | Pod length (cm) | Pod weight (g) | Seed length (mm) | Seed color | Eye color |
| --- | --- | --- | --- | --- | --- | --- | --- | --- | --- |
| **Me51M4-10M5-1M6-M7** | Melakh | Prostrate | 11.5 | 2 | 13.4 | 1.806 | 9.065 | White | Brown |
| **Me51M4-11M5-2M6-M7** | Melakh | Prostrate | 12.2 | 1 | 14.4 | 1.49 | 10.2 | White | Brown |
| **Me51M4-14M5-2M6-M7** | Melakh | Erect | 12.9 | 1 | 16.4 | 2.586 | 12.155 | White | White |
| **Me51M4-25M5-3M6-M7** | Melakh | Semi erect | 10 | 1 | 13.1 | 2.38 | 7.46 | White | White |
| **Me51M4-29M5-1M6-M7** | Melakh | Prostrate | 11 | 1 | 16.2 | 1.912 | 9.315 | White | White |
| **Me51M4-39M5-1M6-M7** | Melakh | Prostrate | 10 | 1 | 15.58 | 2.47 | 10 | White | White |
| **Me51M4-77M5-M7** | Melakh | Erect | 13 | 1 | 6.16 | 1.25 | 16.55 | White | White |
| Me51M4-8M5-3M6-M7 | Melakh | Erect | 10,5 | 1 | 16,6 | 1,73 | 8,685 | White | Brown |
| Me51M4-9M5-1M6-M7 | Melakh | Prostrate | 10 | 2 | 15,8 | 1,76 | 9,035 | White | Brown |
| Me51M4-9M5-3M6-M7 | Melakh | Semi erect | 12 | 1 | 14 | 134 | 7,99 | White | Brown |
| Me51M4-20M5-1M6-M7 | Melakh | Prostrate | 11,3 | 2 | 16,74 | 2,63 | 9,235 | White | White |
| Me51M4-24M5-1M6-M7 | Melakh | Prostrate | 10 | 2 | 19,67 | 2,804 | 11,475 | White | Wite |
| Me51M4-36M5-1M6-M7 | Melakh | Prostrate | 13,2 | 1 | 12,4 | 1,25 | 8,455 | White | White |
| Me51M4-102M5-M7 | Melakh | Erect | 9,33 | 1 | 11 | 0,85 | 12,33 | White | White |
| **Melakh** | IS86-292 x IT83s-742-13 | Semi erect | 12.1 | 1 | 13.4 | 2.423 | 10.25 | White | White |

1 : White ; 2 : White with purple border ; 3 : Light purple ; 4 : Dark purple
Genotypes used for DNA analysis are indicated in bold. For the mutant line codes, the first or second letter refers to the parent's name, the number which
appeared before « M » indicates the selected individual during the M generation.

### Experimental design and development of mutagenized populations

The development of the mutagenized population was performed in different experimental fields located in different sites in the western part of Senegal. In each site, the seeds of each generation were sown using intra row spacing of 50 cm and 75 cm of inter raw spacing. The irradiated seeds (M1) for each variety (Melakh and Yacine) were sown in a separate field in August 2013 around Bambey during the raining season for the development of M1 population. Based on their yield, 12 and 7 mutant plants of Melakh and Yacine respectively were selected and harvested for the development of the M2 population. For these purposes, 103 and 87 seeds respectively from M1 plant of Melakh and Yacine were sown during the dry season in April 2014 in the “Centre National de Recherches Agronomiques (CNRA)” at Bambey (Senegal) to develop M2 population. At maturity, the most productive mutant plants 12 for Melakh and 7 for Yacine were selected, harvested and their seeds sown in the same site (CNRA) in September 2015 to develop the M3 population. From the M3 mutant plants, the most productive were harvested and 88 and 81 seeds for the Mutant Melakh and Yacine were respectively sown in September 2016 in the experimental field located in Ngolgane in the vicinity of Niakhar (Senegal) in accordance with the same experimental design previously described for M2, for the development of M4 population. At maturity stage, the plants were harvested and a single descent method was used to develop M5 population. Thirty-nine (39) and thirty-six (36) seeds of Yacine and Melakh mutants were respectively sown in December 2017 in a pot filled with sand from Sanghalkam (Senegal) and watered with tap water 3 times a week. The mutant plants were grown in the Shadehouse of the “Département de Biologie Végétale” at University Cheikh Anta Diop. The M6 and the M7 population were sown respectively in May 2017 and August 2017 in the field at the Teaching and Research farm of the “Département de Biologie Végétale” at University Cheikh Anta Diop.

### Agro-morphological characterization

Based on the previous studies it is well documented that irradiation promotes the expression of recessive characters in the advanced generations [26, 27]. Therefore, qualitative and quantitative parameters were analyzed from M5 to M7 populations. For instance, in our studies, seed color and pod length variation were noticed in the 5^th^ generation (M5). The scored qualitative parameters encompassed: rate of germination, leaflet abnormalities, leaflet shape, growth habit, flower color, day to first flowering and seed coat color. The quantitative parameters included, percentage of germination, plant height, mean of pod length, number of pod per plant, number of seeds per pod, width and length mean of the seed and weight of 100 seeds. The plant height was measured using a tape measure (Cow head brand) from the cotyledonary node to the top of the plant at the appearance of the first flower [28]. The mean length of 5 pods was measured with a tape measure. Width and length mean of the seed was measured using a vernier caliper (Mutshito) and weighted using a balance (sartoruis®). The data of the quantitative traits recorded throughout M5 to M7 were used for statistical analyses.

### Statistical Analysis of Data

Analysis of variance (ANOVA) and correlation of the quantitative traits were carried out using R software, (R Development Core Team, 2011, version 3.6.1). In order to determine the association between quantitative traits such as the seed mean length, seed mean width, seed mean weight, pod mean length, pod mean width, mean number of seed per pod, plant height and qualitative parameters (like leaflet shape, flower color, growth habit and abnormalities, seed color and flowering date), a standardized Principal Component Analysis (sPCA) was performed with R software using the adjusted means of the measured traits to assess the contribution of each of them on the genetic variability. The phenotypic coefficient of variance, the genotypic coefficient of variation (GCV), the genetic advance (GA) and the broad sense heritability (h^2^) was calculated using R software (R Development Core Team, 2011, version 3.6.1).

The genotypic variance (σ^2^g) was calculated using the following formula:

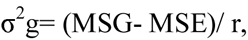

Where MSG is the mean square of genotypes, MSE is the mean square of error, and r is the number of advanced generation.

The Phenotypic Variance (σ^2^p) was assessed as follow:

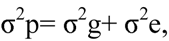

Where σ^2^g is the genotypic variance and σ^2^e is the mean squares of error. Error variance σ^2^e=MSE.

According to [29], the estimation of phenotypic and genotypic coe□cient of variation were calculated as follows:

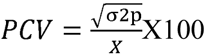

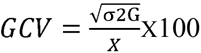, X is the mean.

GCV and PCV values were considered as low (0-10%), moderate (10-20%), and high (□20%) following [30].

The Heritability Estimate (broad sense) (h^2^%), 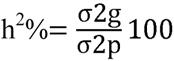

The heritability percentage was considered as low (0-30%), moderate (30-60%), and high (□60%) [31].

The Expected and Estimated Genetic Advance (GA)

GA= k.σp. h^2^, and The Genetic Advance as Percentage of Mean (GA %) is carried using

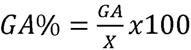

GA was calculated using the method of [32] and selection intensity (k) was assumed to be 5%; where k = 2.06, a constant and σp is the phenotypic standard deviation.

Genetic advance as percentage of mean was categorized as low (0-10%), moderate (10-20%), and high (>20%) [33].

The Pearson’s correlation coefficient (r) for trait linkage evaluation were performed using R software to determine the association between quantitative characters. The genetic distance between mutants and their parents based on quantitative traits was tested using multivariate analysis. To generate the dendrogram, similar matrices were used based on the Ward methods cluster analysis [34].

### DNA genotyping

#### DNA extraction

Five hundred (500) mg of young leaflet collected from each individual plant of 1 month old randomly selected were grounded in mortar in accordance with the protocol developed by Fulton et al. (1995). RNA was removed by adding 50 µg/ml of RNAse A (CalBiochem) in each tube which was incubated at room temperature for 1 h and then DNA was purified according the protocol described by [23]. After precipitation, DNA was dried for 20 min with a Speed Vac ® Plus Sc110 (Savant) and dissolved in 100 µl of TE x 0.1. The quantity and the quality of the DNA extract were performed using a NanoDrop™ One/OneC Microvolume UV-Vis Spectrophotometer (Thermo Scientific) at A260, A280, A260/A280 and A260/A230, then the samples were stored at −20°C or used for amplification.

#### Amplification of DNA and electrophoresis

The amplification reaction was performed in 0.2 ml tube puReTaq Ready-To-Go^TM^ PCR beads (GE Healthcare) containing 2.5 U of lyophilized PuReTaq, 200 µM dNPT and 1.5 mM MgCl_2_, 0.5 µM of each ISSR primer (TSINGKE, China) and 25 ng of DNA in a final volume of 25 µl. The tubes were loaded in a Prime thermocycler (TECHNE, UK) programmed for pre-denaturating step of 5 min at 95°C followed by 40 cycles of 30 s 95°C, 1 min at 38 to 52°C (depending on primer, Table 2), 1 min at 72°C and a final extension of 8 min at 72°C. After amplification, the PCR products were separated on 2% agarose gel (Sigma) for 2 h at 70V. The gel was finally stained during 30 min with GelRed X10,000 (Biotium) according to the manufacturer’s instructions and photographed under UV light using Gel Doc system (High performance UV Transilluminator UVP).

**Table 2.** Estimation of Pearson’s correlation between the agromorphological characters in the M7 of the mutant lines.

|  | PH | Pig | GH | DF | FC | PdL | PdW | SL | SW | SWg | NSP | PdN | PdWg | SC |
| --- | --- | --- | --- | --- | --- | --- | --- | --- | --- | --- | --- | --- | --- | --- |
| PH | 1 |  |  |  |  |  |  |  |  |  |  |  |  |  |
| Pig | 0,31* | 1 |  |  |  |  |  |  |  |  |  |  |  |  |
| GH | 0,72*** | 0,38* | 1 |  |  |  |  |  |  |  |  |  |  |  |
| DF | 0,10 | 0,05 | 0,11 | 1 |  |  |  |  |  |  |  |  |  |  |
| FC | -0,12 | 0,13 | -0,09 | -0,07 | 1 |  |  |  |  |  |  |  |  |  |
| PdL | 0,18 | 0,17 | 0,29 | 0,14 | -0,18 | 1 |  |  |  |  |  |  |  |  |
| PdW | 0,01 | -0,44** | -0,14 | -0,04 | 0,01 | 0,23 | 1 |  |  |  |  |  |  |  |
| SL | -0,09 | -0,26 | -0,19 | -0,27 | -0,10 | 0,13 | 0,46** | 1 |  |  |  |  |  |  |
| SW | 0,11 | 0,15 | 0,22 | -0,19 | -0,17 | -0,04 | -0,47** | 0,02 | 1 |  |  |  |  |  |
| SWg | -0,02 | -0,29 | -0,13 | -0,31* | -0,18 | 0,33* | 0,54*** | **0,74\*\*\*** | 0,27 | 1 |  |  |  |  |
| NSP | 0,31* | 0,18 | 0,36* | 0,11 | -0,20 | **0,82\*\*\*** | 0,26 | 0,10 | 0,04 | 0,31* | 1 |  |  |  |
| PdN | 0,33* | 0,20 | 0,37* | -0,19 | 0,00 | 0,19 | 0,15 | 0,17 | 0,14 | 0,19 | 0,30 | 1 |  |  |
| PdWg | 0,10 | -0,06 | 0,13 | 0,10 | -0,13 | 0,78*** | 0,40** | 0,20 | 0,06 | 0,55*** | 0,67*** | 0,06 | 1 |  |
| SC | -0,35* | **-0,64\*\*\*** | -0,40** | -0,04 | 0,20 | -0,40** | 0,33* | 0,25 | 0,03 | 0,29 | -0,34* | -0,18 | -0,08 | 1 |

\* significant at 5% level of probability; \*\* significant at 1% level of probability; \*\*\* significant at 0.1% level of probability

#### Genetic variation analysis

Amplifications were repeated three times for each single ISSR primer in order to retain the clear and reproducible bands. On this basis, the total number of amplified bands was calculated: their size estimated and the percentage of polymorphic bands evaluated. The polymorphic bands were scored using a binary code as presence (1) or absence (0) to construct a data matrix.

The Shannon diversity index, heterozygosity (Nei index) and the private alleles were calculated using GenAlex 6.5 software [35]. To evaluate the discriminatory power of each marker, the Polymorphic Information Content (PIC) was calculated using the PowerMarker software 3.25 [36], while assessing the genetic variation among and within groups an analysis of molecular variance (AMOVA) was performed using GenAlex 6.5 software [35].

#### Population structure analysis

The population structure was analyzed using the Bayesian clustering approach implemented in the software STRUCTURE 2.3.3 [37] while the number of subpopulations was tested from 1 to 10 independent runs. Using the admixture model [38] each simulation set to 100 000 burn in periods and 10 runs of 200 000 iterations of Markov chain Monte Carlo (MCMC) were performed. These results were uploaded to the online software, Structure Harvester [39] to determine the most likely number of subpopulations using the Evanno Δk method [40]. To assign the individuals into clusters, a membership coefficient q ≥ 0.7 was used. The genotypes within a cluster with membership coefficients q < 0.7 were considered as genetically admixed.

Discriminant Analysis of Principal Components (DAPC) was performed on the basis of the binary matrix data, in order to confirm or invalidate the pattern of the genetic structure obtained with STRUCTURE and to identify the loci responsible for possible differentiation between genetic groups. This analysis was performed using the package « adegenet » [41] of R software [42].

A Neighbor Joining [43] dendrogram was constructed using the inter-individual distance matrix, calculated on the basis of the [44, 45] index. This analysis was performed using Darwin 6.0 software [46].

## Results

### Agromorphological characterization of the munants

#### Variation of qualitative traits among the mutants

The germination rate of the irradiated seeds of Melakh and Yacine was respectively 98.15% and 99.08% in the M1 generation. This germination rate was respectively for Melakh and Yacine 92.23% and 96.5 in the M2, 93.3% and 92.4% in the M3, 63.64 and 81.5% in M4, 90% and 92% in M5, 88% and 93% in M6 and 87.5% and 93.82% in M7 while it was 100% for the control. These results suggested that gamma irradiation at 300 or 340 Gy affected negatively the germinative power of the seeds. Growth habit variability appeared in the M5 for Melakh mutants where 94% were prostrate, 3% erected and 3% semi erected like the control. During the M6, 3% were prostrate, 97% erected and semi erected phenotype was not observed among the mutants. The M7 included 62% prostrate, 7% semi erected and 31% erected. In the Yacine mutants the prostrate phenotype appeared in the M5 with 38% and 61% were erected like the control but in the M6 the percentage of the individuals with these traits was respectively 4% and 96%. The semi erected phenotype appeared for the first time in the M7 with 4% while 18% and 78% were respectively prostrate and erected (Table 1). The leaflet shape within the M7 mutants of Melakh revealed the existence of 4 leaflet forms which were globular (6%), subglobular (76%), hastate (6%) and subhastate (12%) while the leaflet form of the control was subhastate. The leaflet of Yacine was subglobular but 4% of its mutants had subhastate leaflet indicating that gamma irradiation induced leaflet shape variability (Fig 1A-D). Foliar number abnormalities were 11.11%, 11.5% and 0% respectively in M5, M6 and M7 (S1 Fig). These foliar number abnormalities, such as three primary leaves around the node in some mutants instead of two opposite leaves and tetraleaflet, revealed that the gamma irradiation affected the genes controlling leaflet shape in these mutants. Phenotypic variability in flower color was first observed in the M5 of the mutants of Yacine and Melakh. Among the mutants of Melakh 22%, 50% and 36% had white flowers with purple border respectively in M5, M6 and M7 unlike to the control which had white flowers (Fig 1E-H). In contrast, three patterns of flower coloration were observed among the mutants of Yacine. In the M5, 42.5% of the mutant showed white flowers like the control but 35 and 22.5% had white flowers with purple border and purple flowers respectively. The proportion of the mutants with white flowers with purple border was 48% in M6 and 43% in M7 whereas the mutants with purple flowers represented 13 and 26% in the M6 and M7 respectively. These data suggested that gamma irradiation affected the genes controlling flower color of the cowpea samples. Sterility characterized by flower abortion was observed only among the mutants of Melakh in M5, M6 and M7 with 11%, 5% and 2% respectively. Among our population, 50% of flowering was reached 45 days after sowing (DAS) in M6 and M7 for Melakh mutants while this value was respectively 46 and 50 DAS for the mutants of Yacine (S2 Fig). The color of the seed coat was unchanged from M1 to M7 for the mutants of Melakh but some of them showed brown or beige eye. In the M4 of Yacine mutants, the seed coat was brown like the control except one mutant where brown seeds, white seeds and pigmented brown and white were harvest (Fig 1J). In the M5, 43% had white seeds, 38% of the mutants had brown seeds as the control and 19% had light brown seeds (Fig 1K-M).

**Fig 1.**
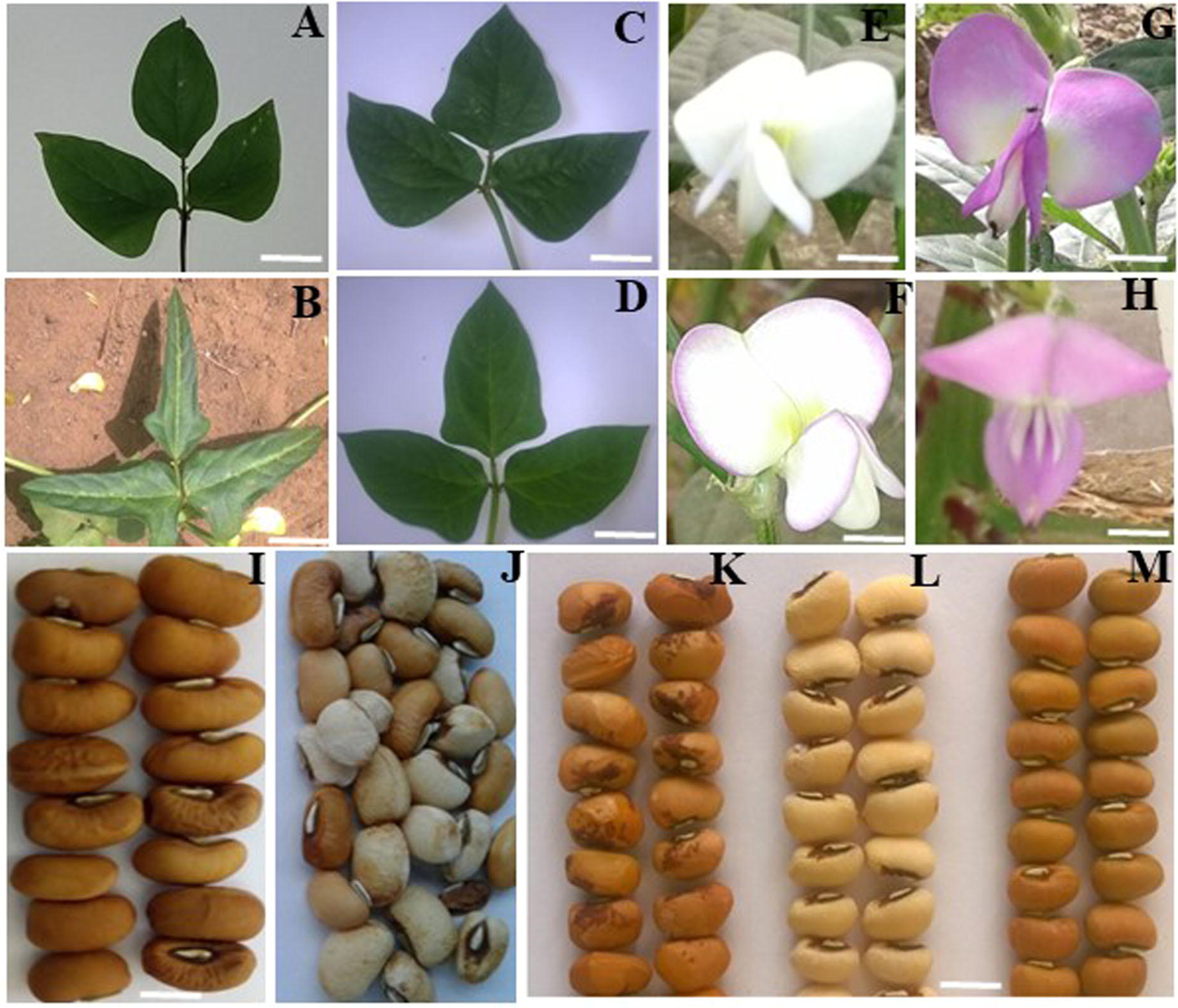
Variation of qualitative traits observed in the populations. **A**-**D**: Leaflet shape, Bar = 6.25 cm; **E**-**H**: Flower color, Bar = 0.84 cm; **I**-**M**: Seed coat color, Bar = 1 cm **A**: Globular (Yacine); **B**: Hastate; **C**: Subglobular; **D**: Suhastate (Melakh); **E**: White flowers (Yacine and Melakh), **F**: White flower with purple border; **G**: Dark purple; **H**: Light purple; **I**: Brown seed coat (Yacine); **J**: Brown, white, white pickleed brown seed coat (from on single plant in M4 of the mutant of Yacine); **K**: light brown speckled in dark brown seed coat; **L**: White seed coat; **M**: Light brown seed coat

#### Variation of quantitative traits and yield components among the mutants

To advance the mutant populations from M1 to M4, pedigree method was used and the selection criteria was based on the plant yield and no shattering pods for the mutants of Yacine only. In contrast, from M5 to M7, single seed descent method was used. In M5, the mutants of Melakh were selected based on 100 seed weights but in M6 and M7 one more yield component (pod length) was including in the selection criteria. The average pod length of Melakh control measured across generations was 19 cm while the value obtained in the mutants ranged from 12.5 to 25 cm cross generation M5, M6 and M7 (Fig 2A). At the same time, the average pod length of Yacine control was estimated to 14.65 cm but variability of the pod length was observed among the mutants of Yacine ranging from 8.70 to 19 cm cross generations M5, M6 and M7.

**Fig 2.**
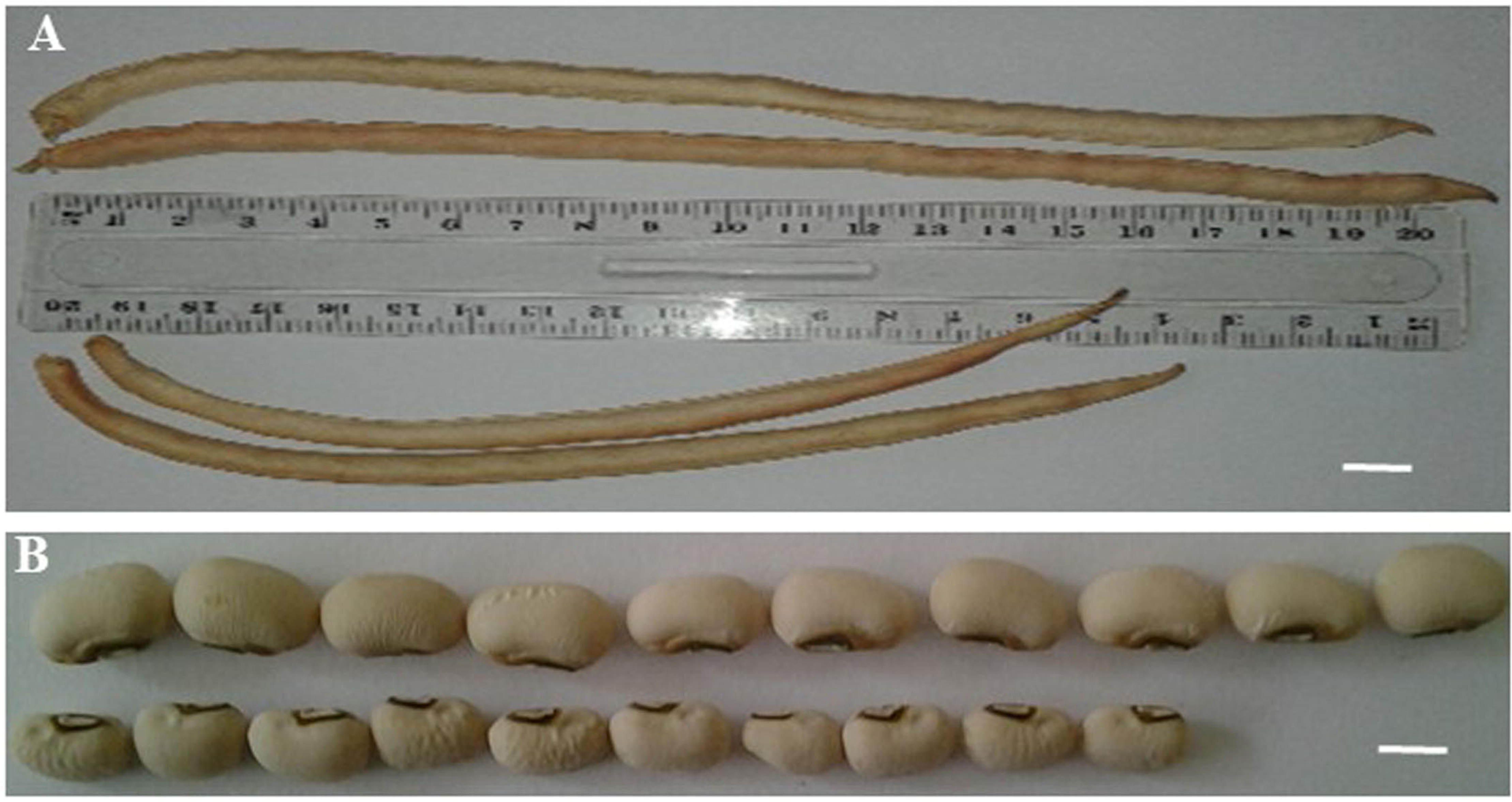
Variation of the pod length (**A**) and seed size (**B**) in M5 among the mutants of Melakh. On the top: Pods and seeds harvested from the mutant Me51M4-39M5. Bar = 1.54 cm On the bottom: Pods and seeds harvested from the parent control Melakh. Bar = 1 cm

The number of pods per plant ranged from 1 to 6, 1 to 20 and 2 to 15 in M5, M6 and M7 respectively for the mutants of Yacine, for the control it ranged from 3 to 6. These values ranged from 1 to 7, 2 to 35 and 1 to 43 for the mutants of Melakh and from 3 to 10 for the control. The mean seed length varied from 8.75 to 10.72, 8.1 to 11.66, 7.46 to 12.16 mm respectively for M5, M6 and M7 for the mutants of Melakh and 9.2 mm for the control (Fig 2B). For the mutants of Yacine the mean seed length varied from 6.12 to 10.04, 9.12 to 11.20 and 8.11 to 11.13 mm respectively for M5, M6 and M7 and 10 mm for the control. The number of seeds per pod varied from 9 to 15, 4 to 12 and 3 to 12 for the mutants of Melakh respectively for M5, M6 and M7 and 10 for the control. For the mutants of Yacine, the number of seeds per pod ranged from 7 to 13, 5 to 12 and 4 to 16 respectively in M5, M6 and M7 and 9 for the control. The 100 seed-weight ranged from 16.67 to 28.52, 20.07 to 32.28 and 13.55 to 38 g respectively for M5, M6 and M7 for the mutants of Melakh but the values recorded for the control ranged from 18.35 to 32.12 g. At the same time, 100 seed-weight ranged from 10.78 to 24.5, 17.27 to 33.16, 13.83 to 30.03 g respectively in M5, M6 and M7 population of the mutants of Yacine and from 16.4 to 30.40 g for the control. Two mutant lines of Melakh produced more seeds (44 to 184) per plant regardless of the generation. Similar results were observed among the mutants of Yacine (17 to 150 seeds).

#### Genotypes clustering based on Principal Components Analysis and correlation between traits

The projection of agromorphological parameters collected from M7 in the PCA Biplot showed that the axis 1 explained 28% of the variation. This axis encompassed the elite mutants in term of seed length, pod weight and seed weight (Y7-M4-3M5-1M6-M7 and Me51M4-14M5-1M6-M7) and the early maturing mutant lines (Me51M4-36M5-1M6-M7, Y1-M4-16M5-2M6-M7 and Y7-M4-1M5-1M6-M7) compared to their control parents Yacine and Melakh. The axis 2 explaining 19.7% of the variation, was constituted by the mutant lines (Y1-M4-11M5-3M6-M7, Y7-M4-1M5-3M6-M7, Y1-M411M5-3M6-M7, Me51M4-39M5-1M6-M7, Me51M4-29M5-1M6-M7, Me51M4-20M5-1M6-M7, Me51M4-10M5-1M6-M7, Me51M4-9M5-1M6-M7, Me51M4-11M5-1M6-M7) which acquired a new growth habit i.e prostrate compared to their parents (Fig 3). The evaluation of the Pearson’s coefficient between agromorphological characters cross generation (from M5 to M7) showed that steam pigmentation (Pg) was significantly and negatively correlated to seed color (SC) (r = −0.5, p = 0.01 in M5; r = −0.59, p = 0.001 in M6 and r = −0.64, p = 0.001 in M7), the number of seeds per plant (NSP) was significantly and positively correlated to the pod length (PdL) (r = 0.42 p = 0.05 in M5, r = 0.62 p = 0.001 in M6 and r = 0.82 p = 0.001 in M7), Seed weight (SWg) and Seed Length (SL) (r = 0.86 p = 0.001 in M5, r = 0.60 p = 0.001 in M6 and r = 0.74 p = 0.001 in M7) (S1 Table, S2 Table and Table 2). In addition to these, in the M7, which is supposed to be more stable, the yield component parameters such as the pod number (PdN) and the growth habit (GH) are significantly correlated (r = 0.37 p = 0.05) and pod weight (PWg) and PdL (r = 0.78 p = 0.001). The pod width is significantly and positively correlated to the SWg (r = 0.54 p<0.001), PWg (r = 0.40.01 to SL (r = 0.46 p = 0.01) as do the pod weight (PdWg) and SWg (r = 0.56 p = 0.001). In contrast, Seed width (SW) and PdW are negatively and significantly correlated (r= −0.47 p = 0.01, Table 2).

**Fig 3.**
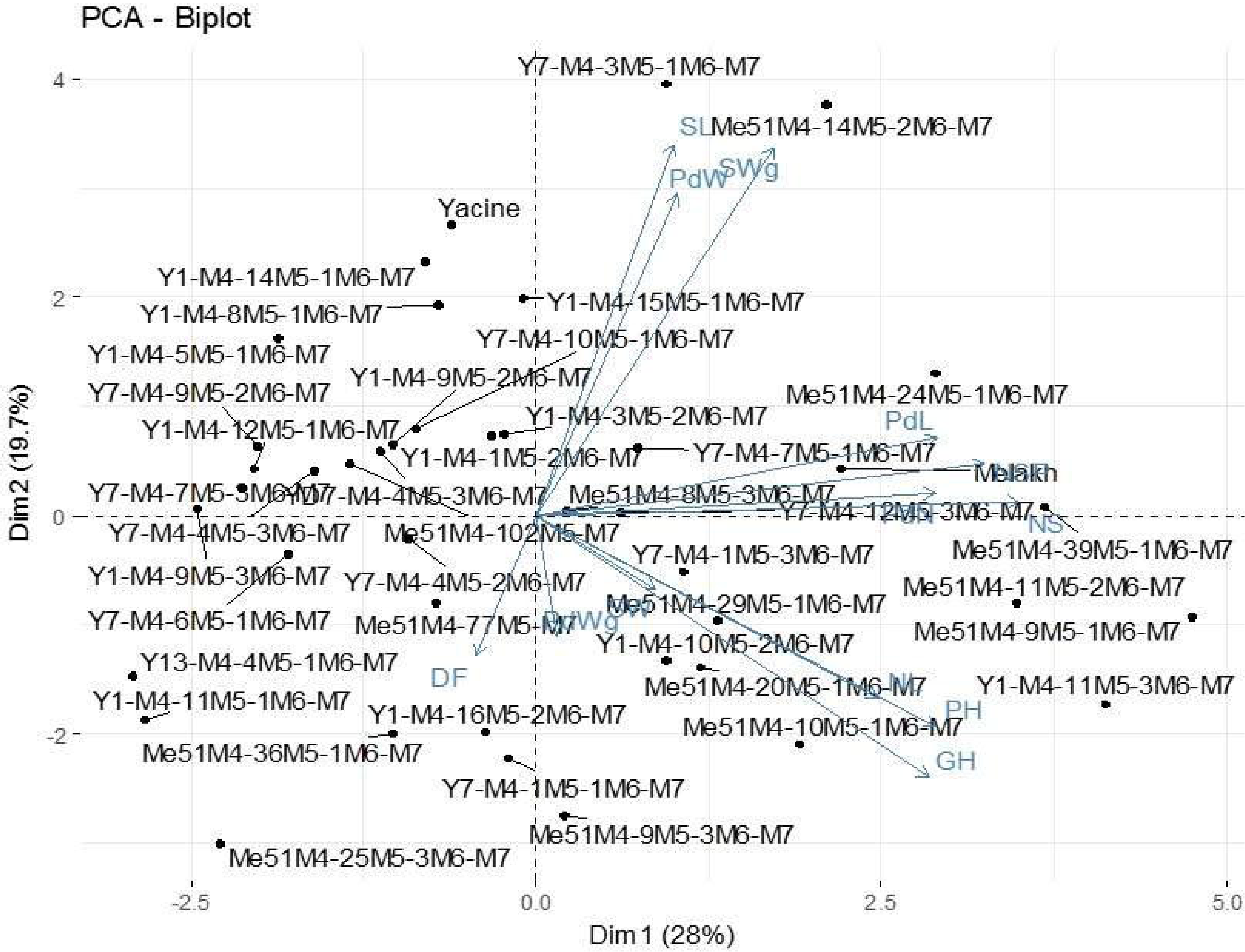
Principal Component Analysis of 40 mutant M7 lines and their parent based on agro-morphological data.

#### Relationship among mutants based on agromorphological parameters

Based on the agro-morphological characters the M7 population was divided into 3 groups. The first group which encompassed 52% of the individuals are divided into 3 subgroups. The subgroup 1 included only the mutants (Me51M4-102M5-M7, Me51M4-36M5-1M6-M7, and Me51M4-77M5-M7) which are sister of Y13M4-4M5-1M6-M7, Y1M4-11M5-1M6-M7 and Me51M4-25M5-3M6-M7. The subgroup 2 only comprised mutants of Yacine except Me51M4-8M5-3M6-M7 which clustered with genotypes sharing the same characters such as erected port, white brown-eyed seed. The subgroup 3 included only mutants of Yacine. The second group included 14% of the genotypes which had all erected port and brown-eyed seed and contained only mutants of Yacine and their parent. The third group was the second largest one with 33% of the genotypes was divisible into 2 main subgroups. The first subgroup encompassed Melakh its mutants and 2 mutants of Yacine. The second included mutants of Melakh and 2 mutants of Yacine which shared several agromorphological characters with Melakh among these long pod and white seed (Fig 4).

**Fig 4.**
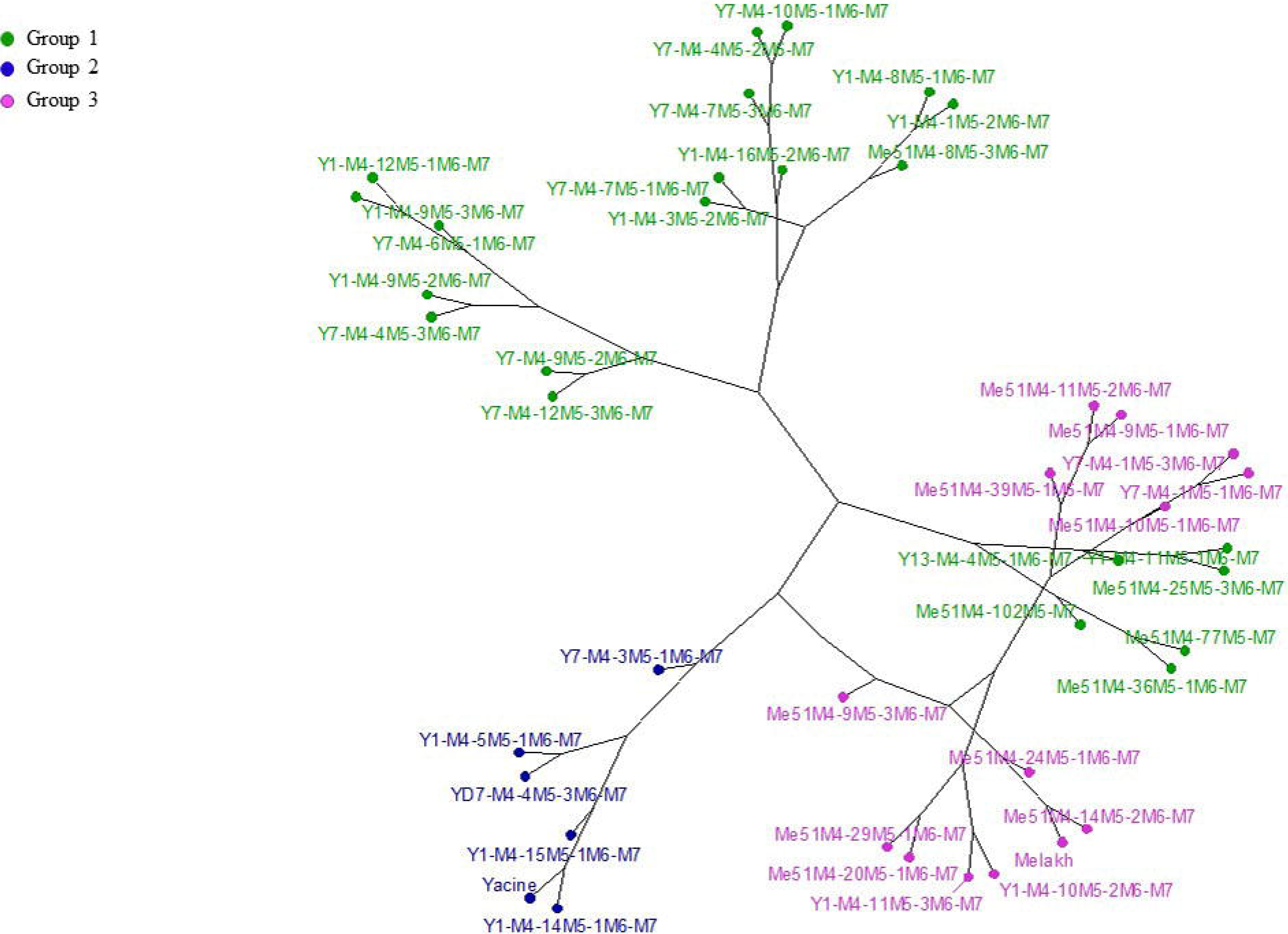
Hierarchical classification of 40 mutant lines and their parent based on agro-morphological characters using Ward’s method in R software.

#### Heritability of agromorphological characters in the mutants

Using R software, it was possible to analyze the phenotypic (PCV) and genotypic (GCV) coefficient of variance and the heritability values for the different measured traits in the mutant populations. The PCV varied from 100.59 for pod number (PdN) to 9.12 for Seed length (SL). Plant height (PH) recorded high value (75.1%) of PCV in contrast, day of flowering (DF), pod width (PdW), eye color (HC) and leaflet length (LF) showed respectively 9.37, 10.14, 10.32 and 15.89%. The recorded GCV values ranged from 67.12% for pod number (PdN) to 5.97% for date of flowering (DF). All the studied traits showed a genetic coefficient of variance (GCV) below 50% except pod number and seed color (SC). The genetic advance (GA) as a percentage of the mean recorded in the mutants ranged from 0.45% for petiole length to 92.27% for pod number. High values of heritability were recorded for seed color (94.48%), steam pigmentation (92.83%), flower color (78.01%), eye color (68.98%), pod length (68.29%), pod width (59.06%), seed weight (57.58%), number of seed per plant (52.11%), seed length (51.19%), the remaining characters showed values below 50% (Table 3).

**Table 3.**
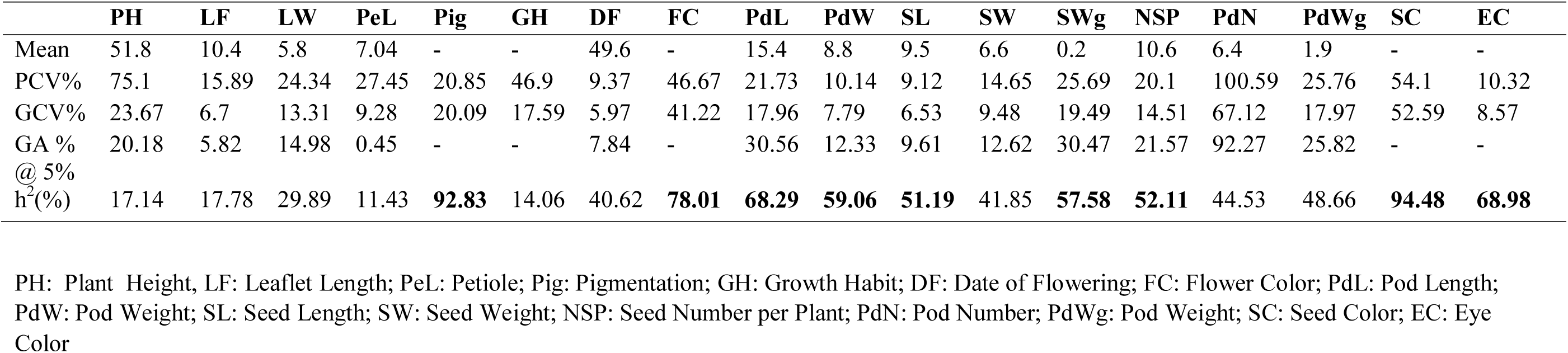
Estimation of mean values of phenotypic coefficient of variance (PCV), genotypic coefficient of variation (GCV %), broad sense heritability (*h2bs*%) and genetic advance as% of the mean (GA %) of eighteen traits (quantitative and qualitative) in the M7 of 40 cowpea mutant lines.

### Genetic characterization of the mutants

#### Genetic diversity induced by mutagenesis revealed by ISSR markers

In total, the 13 ISSR markers used in this study gave polymorphism amplifying 129 bands (loci) in the 18 mutant lines and their 2 parents. The size of the amplified bands ranged from 150 to 2000 bp. Of these, 111 (86%) bands were polymorphics with numbers ranging from 5 (UBC827) to 12 (UBC825 and UBC844) for each primer (Table 4). The percentage of the polymorphic bands per primer ranged from 60% (UBC809) to 100% (UBC825, UBC844, 17899A, 17899B and, HB10). The band frequencies ranged from 0.318 (UBC841) to 0.808 (HB12) with a mean of 0.473±0.145. The genetic diversity ranged from 0.142 (UBC841 and UBC823) to 0.293 (UBC844 and 17899B) with a mean of 0.220±0.049. The Polymorphic Information Content (PIC) values for each primer varied between 0.167 (UBC841) and 0.307 (HB09) with a mean of 0.238±0.044 (Table 5). Based on these data, our analyses showed that the average diversity level for all the mutants and their parents was equal to 0.222 (Nei index). The level of the genetic diversity observed in group 1 (h = 0.308) was higher than the one in group 2 (h = 0.146; p = 0.0001) and in group 3 (h = 0.212; p = 0.0001). The level of genetic diversity in group 3 was higher than in group 2 (p = 0.016). The genotypes which belong to group 1 also recorded a greater number of private alleles (17 bands vs. 4 for group 2 and 7 for group 3, Table 6). The Shannon’s information index was higher in group 1 (0.463) than in group 2 and 3 with respectively 0.211 and 0.319 but for the entire population this value was 0.331. To investigate the genetic variance within and among genetic pools, AMOVA was carried out in this study. Our results revealed that the majority of the variance rather existed within group (85%) than among group (15%) (Table 7).

**Table 4.** List of the Inter-simple sequence repeat (ISSR) primers with their annealing temperatures (Tm) used to genotype the mutants M7 and their parent, total number of bands, number of polymorphic bands, percentage of polymorphic bands and the band size range.

| ISSR primer codes | Sequences (5'-3') | Annealing temperature (°C) | Total number of bands | Number. of polymorphic bands | Percentage of polymorphic bands (%) | Band size range (pb) |
| --- | --- | --- | --- | --- | --- | --- |
| UBC809 | (AG)8G | 52.2 | 10 | 6 | 60 | 1400-200 |
| UBC825 | (AC)8T | 49.8 | 12 | 12 | 100 | 1500-200 |
| UBC841 | (GA)8YC | 52.6 | 14 | 11 | 78.57 | 1800-150 |
| UBC844 | (CT)8AC | 52.6 | 12 | 12 | 100 | 1300-300 |
| 17899A | (CA)7AG | 49.2 | 11 | 11 | 100 | 2000-250 |
| 17899B | (CA)7GG | 51.7 | 11 | 11 | 100 | 1500-300 |
| HB8 | (GA)6GG | 44 | 8 | 6 | 75 | 1300-250 |
| HB9 | (GT)6GG | 44 | 10 | 8 | 80 | 1200-250 |
| HB10 | (GA)6CC | 44 | 8 | 8 | 100 | 1000-350 |
| HB12 | (CAC)3GC | 38 | 7 | 6 | 85.71 | 1500-300 |
| UBC827 | (AC)8G | 52.2 | 7 | 5 | 71.42 | 1000-400 |
| UBC823 | (TC)8C | 52.2 | 10 | 8 | 80 | 1500-400 |
| UBC807 | (AG)8T | 49.8 | 9 | 7 | 77.77 | 1500-200 |

**Table 5:** Band frequency, genetic diversity and polymorphism information content (PIC) of each ISSR locus.

| Marker | Band Frequency | Gene diversity | PIC |
| --- | --- | --- | --- |
| ISSR 825 | 0.338 | 0.203 | 0.219 |
| ISSR841 | 0.318 | 0.142 | 0.167 |
| ISSR823 | 0.381 | 0.142 | 0.172 |
| ISSRHB-10 | 0.369 | 0.232 | 0.228 |
| ISSRHB-8 | 0.575 | 0.231 | 0.236 |
| ISSR17899A | 0.468 | 0.212 | 0.224 |
| ISSRUBC-807 | 0.55 | 0.228 | 0.242 |
| ISSR827 | 0.46 | 0.236 | 0.293 |
| ISSR844 | 0.417 | 0.293 | 0.288 |
| ISSRHB-09 | 0.413 | 0.274 | 0.307 |
| ISSR17899B | 0.368 | 0.293 | 0.274 |
| ISSR HB-12 | 0.808 | 0.164 | 0.196 |
| ISSR809 | 0.683 | 0.211 | 0.258 |
| Mean | 0.473±0.145 | 0.220±0.049 | 0.238±0.044 |

**Table 6.** Statistical analyses of genetic diversity level.

| Population | Number of<br>genotypes | % of<br>polymorphic<br>loci | Nei diversity | Shannon's<br>Information<br>Index |
| --- | --- | --- | --- | --- |
| Group1 | 7 | 85.59 | 0.308 | 0.463 |
| Group2 | 4 | 34.23 | 0.146 | 0.211 |
| Group3 | 9 | 61.26 | 0.212 | 0.319 |
| All genotypes | 20 | 60.36 | 0.222 | 0.331 |

**Table 7.** Genetic variance within and among groups based on Analysis of Molecular Variance (AMOVA).

| Source of variance | Degree of freedom | Sum of Square | Mean Square | Estimated Variance | Percentage of total variance |
| --- | --- | --- | --- | --- | --- |
| Among group | 2 | 65.008 | 32.504 | 2.729 | 15 |
| Within group | 17 | 257.992 | 15.176 | 15.176 | 85 |
| Total | 19 | 323.000 |  | 17.905 | 100 |

#### Population structure and genetic relationship among genotypes

Using STRUCTURE software [37], the evaluation of the delta k according to the Evanno method showed the highest peak at k = 3 (Fig 5A) and the mean value of the logarithm of likelihood (LnP (D) for k = 1 was lower than that of k = 3 which is the higher peak (Fig 5B). The representation of the ancestry at k = 3, revealed three (3) genetic pools (Fig 5C). Group I included 30% of the individuals while group II and group III encompassed respectively 50 and 20% of the genotypes. The genetic relationship revealed by the dendrogram was in agreement with STRUCTURE analysis which clearly distinguished three groups. The first group contained 2 mutants of Melakh (Me51M4-14M5-2M6-M7 and Me51M4-39M5-1M6-M7) and 4 other mutants of Yacine (Y1-M4-8M5-1M6-M7, Y1-M4-3M5-2M6-M7, Y1-M4-11M5-1M6-M7 and Y7-M4-4M5-3M6-M7). This group encompassed the highest number of admixed individuals. The second genetic pool contained exclusively the mutants of Yacine. Of these, Y1-M4-9M5-2M6-M7 and Y1-M4-9M5-3M6-M7clustering with a high bootstrap value (96%). The admixed Y7-M4-1M5-1M6-M7 clustered with Y7-M4-6M5-1M6-M7 with 80% bootstrap value. The third genetic pool contained Yacine, 3 mutants of Yacine (Y13-M4-4M5-1M6-M7, Y7-M4-12M5-3M6-M7 and Y7-M4-10M5-1M6-M7) and 5 other mutants of Melakh (Me51M4-10M5-1M6-M7, Me51M4-11M5-2M6-M7, Me51M4-25M5-3M6-M7 and Me51M4-29M5-1M6-M7). The variety Melakh was clustering with its mutant, Me51M4-77M5-M7 with a high bootstrap value (92%) (Fig 6A).

**Fig 5.**
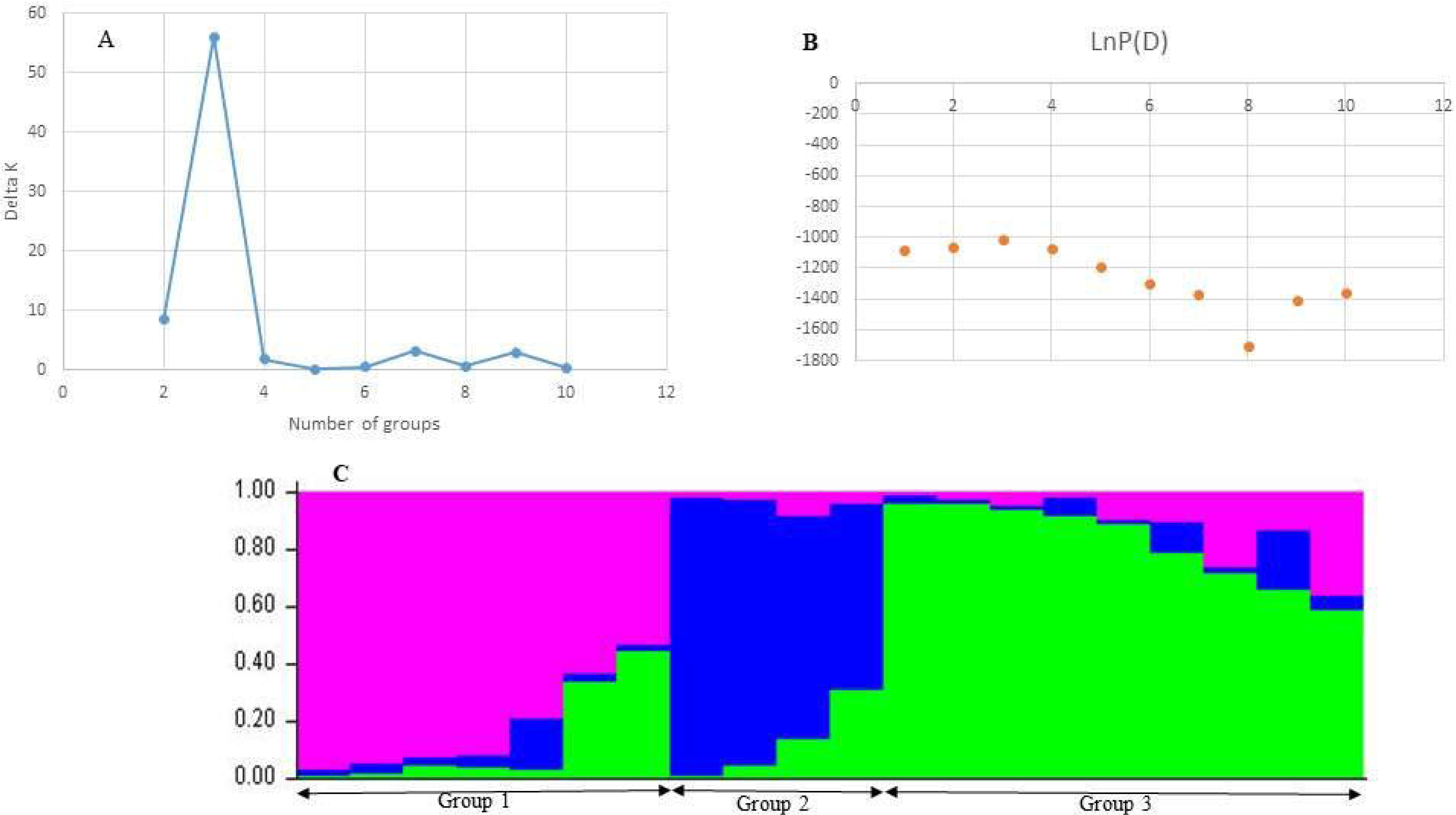
Structure of the mutant population. **A**: Probability of subdivision into genetic groups by the method of Evanno et al. 2005; **B**: Probability of subdivision into genetic groups by the log mean likelihood method and **C**: Ancestry for K = 3.

**Fig 6.**
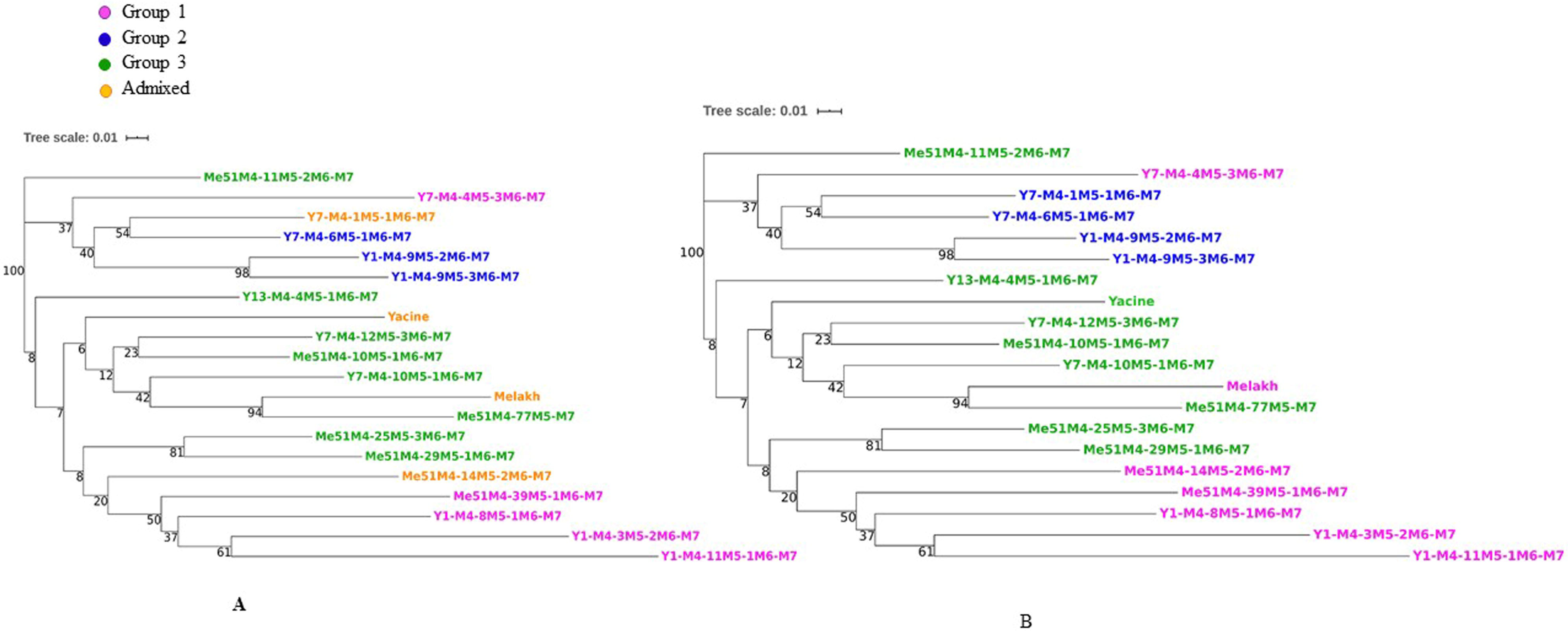
Neighbor Joining Dendrogram based on ISSR data representing the grouping of 18 cowpea mutant lines and their parent. The dendrogram was constructed using the distance matrix between individuals, calculated using the Jaccard, (1902, 1912) Index. **A** and **B**: Dendrograms based on STRUCTURE and DAPC analyses respectively.

The clustering of the genotypes resulting from the DAPC analysis identified 3 groups which showed any individual genetically admixed (Fig 6B). This results suggested that DAPC analysis was appropriate to assess mutant populations structure by achieving better separation among group as it was also showed by the number of clusters identified (Fig 7A). Group 1 and Group3 differed from each other based on the first axis of the DAPC. This classification was based on the loci 84, 37, 57, 105, 35, 10 and 106 which were the most discriminating, in decreasing order (Fig 7B). Group 2 differed from the others on the second axis. These findings were based on the loci 97, 78, 87, 21, 105, 7, 82, 64, 47 and 12 which were the most discriminating, in decreasing order (Fig 7C).

**Fig 7.**
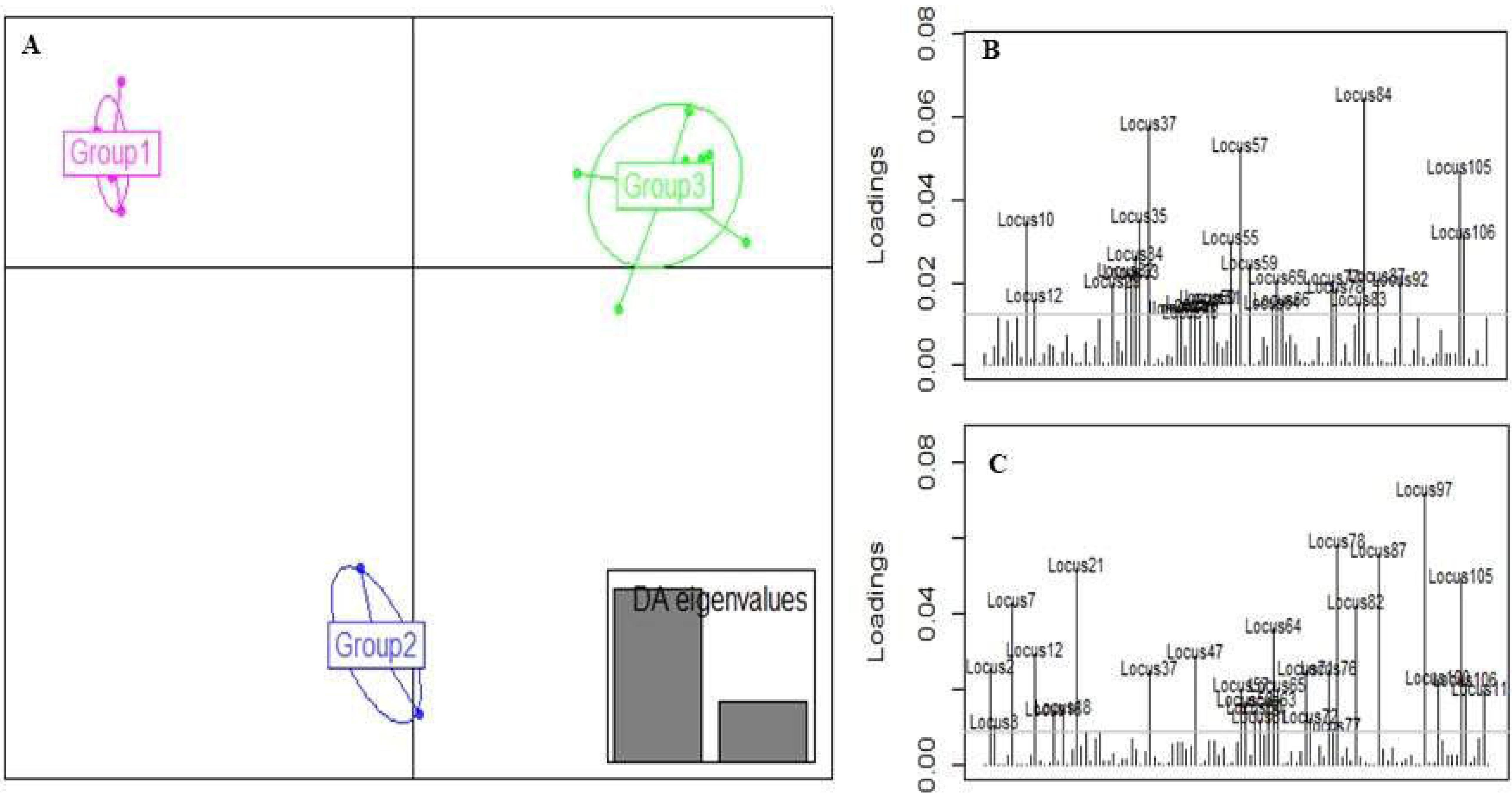
Discriminant Analysis of Principal Components (DAPC) for 18 cowpea mutant lines and their parent based on ISSR data. **A**: graphical representation of the groups; **B**: contribution of the alleles based on the first axis; **C**: contribution of the alleles based on the second axis

## Discussion

Wide genetic variability is a prerequisite for a successful breeding program particularly in the era of climate change with its adverse effects leading to the erosion of the plant genetic resources. Thus, to broaden the crop’s genetic basis, a wide range of techniques have been used in the last decade but among these, induced mutagenesis has been proved to be best for creating novel variation in crop genome and was used in this study to expand variability in cowpea which experienced a single domestication event during the course of evolution.

### Agromorphological variability analysis of the mutants

In this study the percentage of germination of the irradiated seeds decreased compared to the control (Melakh and Yacine) regardless of the dose and the generation used. These results were in agreement with the findings of Melki and Marouani [47], Horn et al. [13] and Olasupo et al. [17] who recorded similar observations during their studies. In contrast, Horn et al. [13] recorded zero germination of cowpea irradiated seeds at 300 Gy, in this study 98.15% of field establishment were observed for Melakh M1 white-seeded at this dose, this value reached 99.08% for Yacine M1 brown-seeded at 340 Gy. These results were in agreements with the findings of Olasupo et al. [17] who suggested that radio-sensitivity is genotype dependent as it was associated with seed characteristics (seed coat color, water content, thickness and weight). Presently, it is well documented that ionizing radiation is injurious of enzymes and growth hormone leading to biochemical and physiological modifications, cell death, abnormal cell division, tissue and organ failure and growth disturbance [48, 49, 50]). We can assume that these type of changes occurred in our irradiated seed materials as 11.11% and 11.5% of the Melakh mutant lines in M5 and M6 respectively showed abnormities of leave numbers, corroborating the discoveries of Girija and Dhanavel [51] and Nair and Mehti [11] performed in cowpea mutants that were bifoliated, tetrafoliated, pentafoliated and hexafoliated (S1 Fig). These disturbances would explain the change observed in leaves shape of our mutants with 3 new forms e.i. globular, subglobular and hastate (Fig 1). In contrast, the investigations performed by Gnanamurthy et al. [16] led to the discovery of only a globular form within their mutant populations. Similar modifications induced early flowering in some Melakh and Yacine mutant lines which is an important agronomic trait for farmers and breeders particularly in the Sahel zone where the rainy season has become shorter and shorter. In addition, mutagenesis treatment affected flower coloration which changed from white to white with purple border in Melakh and in Yacine mutant population it changed from white to white with purple border, purple or dark purple with a high heritable value (*h*^2^ = 78.01%) (Fig 1, Table 3). These findings were in accordance to the observations of Horn et al. [13] and Girija et al. [52] who also noticed flower color variation in their cowpea mutant populations. In this study, similar seed coat color variation was observed in the mutants of Yacine, first time during the M4 generation where one single plant produced white seeds, brown seeds and white-browned seed (Fig 1J) but from M5 brown seed and white were only recorded as it was previously reported that seed coat color is an important trait for consumer preference depending on the regions [53]. On the other hand, the seed coat color of Melakh mutants remained unchanged e.g white as the control regardless of the generation. These results explained the high heritability value (*h*^2^ = 94.48%; Table 3) recorded for seed coat color in our populations. In accordance with this study, mutants seed coat color variation was also recorded by Gaafar et al. [12] and Horn et al. [13] suggesting that mutation affected the candidate genes involved in the control of late stages in flavonoid biosynthesis pathway namely the basic helix–loop–helix gene for the *C* locus, the WD-repeat gene for the *W* locus and the *E3 ubiquitin ligase* gene for the *H* locus [54].

During this study, yield component (pod length, pod width, number of seeds per plant, seed length and seeds weight) variation with a high heritability values was noticed among the mutant genotypes compared to the control. Similar results were recorded by Gnanamurthy et al. [16] Goyal and Khan [55] and Horn et al. [13] which suggested that mutagenesis can be used to improve crop yield, one of the most important agronomic characteristics for breeders. High heritability, genetic advance and genetic coefficient of variation values of pod length and number of seeds per plant recorded in this study suggest that these traits can be considered as attributes for the improvement through selection of the mutants. In addition, regardless of the generation (M5 to M7) and the environment, the pod length and the number of seeds per plant were positively and significantly correlated as did seed length and seed weight during this study. In contrast, the variation of the correlation values between traits recorded might be due to the effect of interaction genotype x environment which could induce epigenetic modifications impacting gene expression or gene segregation over generations.

The genetic relationship based on the agromorphological characters revealed that the mutant populations were subdivided into 3 groups (Fig 4). The first group included genotypes which notably deviated from their respective parents, thereby suggesting that the irradiation doses used were efficient enough to induce heterogeneous population as any polytony was observed in the dendrogram. In the remaining two others groups, most of the mutants were clustering with their respective control revealing their high phenological diversity as reported by Laskar and Khan [56]. Taking together, these findings meet the recommendation of Rohman et al. [57] who suggested that the cluster contributing to the greatest divergence can help to choose the parent in a breeding program. Based on these, PCA results (Fig 3) revealed in this study that yield related traits such as pod length, number of seeds per plant and seed weight are major contributor to the genetic divergence which was in accordance with the report of Afuape et al. [58] who suggested that PCA is accurate to select the best genotypes for future breeding progam.

### Genetic diversity of mutants based on molecular markers

In the present study, the gene diversity measures through the average heterozygosity and the genetic distance among individuals in a population and the PIC values which is a good indicator of the usefulness of a marker to determine its inheritance between the offspring and the parent were estimated in our mutants. Our results showed that, in general, the gene diversity and the PIC were close (Fig 5), suggesting the evenness of allele’s frequencies in the mutant populations as reported by Shete et al. [59]. According to Botstein et al. [60], a marker is considered highly informative if the PIC is ≥ 0.50, moderately informative with a PIC values ranging from 0.25 to 0.5 and slightly informative at less than 0.25. Based on this, the ISSR827, ISSR844, ISSRHB9, ISSR17899B and ISSR809 were moderately informative, the remaining markers were slightly informative which were higher than the scores recorded in the cowpea mutant population developed by Gaafar et al. [12]. These differences could be attributed to the low irradiation dose (50 Gy) used which might induce less mutations and less variability compared to the 300 Gy and 340 Gy employed in this study. In addition, Gaafar and coworkers [12] used ethidium bromide for DNA staining which is less sensitive than GelRed, the one we used in our study leading to more DNA bands scoring. In contrast, the PIC scores reported in the natural populations of *Vigna* [61], mungbean, blackgram [62] and in *Vigna unguiculata* [21] were a bit higher probably due to several mutations undergone by these genomes during evolution. Our results show that ISSR are accurate markers to discriminate cowpea mutants for new genotypes identification and selection. In this study, the ISSR primers UBC825 (AC repeat motif), UBC844 (CT repeat motif), 17899A and 17899B (CA repeat motif) and HB12 (GA repeat motif) gave 100% polymorphic bands which suggested that these motifs are abundant in the genome of cowpea. Similar observations have been reported in *Arachis hypogea* [63, 64] and in *Vigna mungo* [65].

#### Population structure and genetic relationship of the mutants based on ISSR markers

Analyzing population structure in mutants is relevant to understand the organization of the genetic variation which is driven by the combined effect of recombination, mutation, demographic history and natural selection. Based on this, and due to the informative nature of the ISSR markers used, the generated data subjected to STRUCTURE analysis showed 3 subpopulations (optimal K = 3). These results were consistent with the organization of the dendrogram (Fig 6A). The clustering of the Me51M4-39M5-1M6-M7 and Y1-M4-8M5-1M6-M7, two genotypes tolerant to the nematode *Meloidogyne incognita*, compared to their parents, according to our preliminary studies, might suggest that this character is distributed in this group (group 1). In addition, this group encompassed genotypes with long pod length compared to their respective parents. These findings are relevant to select the best mutant genotypes which can be proposed for variety registration and popularization but also for gene discoveries and breeding programs. These results demonstrate the ability of gamma rays to induce large genetic changes in DNA material as among the 18 mutants genotyped 7 derived from 1 Melakh mutant and 11 from 1 Yacine mutant. The findings were in accordance with previous studies recently performed on several crops such as cowpea [12,13,66] and chickpea [67]. The dendrogram analysis revealed a group encompassing only Yacine mutants. In this group 2, the Y7-M4-4M5-3M6-M7 genotype could be included as it shares several morphological characters such as the erected shape, except Y7-M4-1M5-1M6-M7 which is prostate, white flower with purple border except the mutant Y1-M4-9M5-2M6-M7 showing light purple flower, brown coat seed, except Y7-M4-1M5-1M6-M7 with white coat seed, Y7-M4-6M5-1M6-M7 a white-brown seeded genotype as is Y1-M4-9M5-3M6-M7. Thus, ISSR are accurate markers to discriminate these genetic lines. In contrast, the group 3 is more heterogeneous including Yacine, its mutants and its progenitor Melakh and its mutants. Those results suggested that the assignation of Melakh to the group could be an artefact. The clustering of Yacine and its relative Melakh in the same group 1 demonstrate the discriminating power of the DNA based molecular markers.

#### Genetic differentiation within and between groups

In the present study, STRUCTURE analysis and the dendrogram results showed 3 groups which aroused our curiosity to understand the variability within and between clusters by assessing the genetic parameters such as Nei’s genetic diversity and Shannon’s information index which are important to measure the degree of genetic diversity among and within group in a population. Our results showed that group 1 had a high Nei’s genetic diversity and Shannon’s information index and a high number (17) of private alleles unlike group 2 which had the lowest diversity, suggesting that the gamma irradiation doses were efficient to induce new alleles in the mutants (Table 6). Identification of private alleles are important in plant breeding and conservation as their presence in a single population might be linked to specific agronomic traits usable for new genotypes selection in a mutant population. In accordance with this, the AMOVA results revealed that a large majority of the total variation (85%) was noticed within group variation suggesting a high level of differentiation while only 15% of the variation were recorded between group (Table 7). According to Seyoum et al. [68], small variation between groups might be an advantage due to its usefulness to study marker-traits association. In contrast, a large variation between groups could reduce the possibility to detect the effects of single genes in a genome wide association study [69]. Based on these and taking into account the important agronomic traits recorded in our mutant populations a genome wide association can be performed in order to detect the molecular markers associated with these characters and usable in a molecular breeding program.

To analyze population structure, different methods such as STRUCTURE, Principal Coordinate Analysis and DAPC can be used. The latter provides complementary information leading to better assignments of individuals in the accurate group and its advantage is related to the fact that the target population has not to be in Hardy-Weinberg equilibrium. In this study, DAPC results divided the mutant populations in well-defined 3 groups with less admixture compared to STRUCTURE results. Indeed, the ancestry value recorded in STRUCURE analysis allowing the assignment Me51M4-14M5-2M6-M7 and Melakh to group 1 was less than 70% in contrast, this abnormality was resolved using DAPC as it did with Y7-M4-1M5-1M6-M7 and Yacine to group 2 and 3 respectively. These results were in accordance with previous studies carried out in potato [70], in landraces and bred cultivars of *Prunus avium* [71] and Ginseng germplasm [72] which revealed that DAPC gave a better grouping resolution than STRUCTURE. In addition, DAPC analysis showed that the loci such as 35 and 37 (amplified by ISSR primer HB10), 57 (amplified by ISSR primer 807) greatly contributed for the discrimination of group 1 and group 3. In contrast loci 97, 78, 21 and 105 amplified respectively by 17899B, ISSR844, ISSR 841 and ISSRHB12 was also involved in the differentiation of group 2 from the others (Fig 7B). This latter ISSR primer was suspected by Gaafar et al. [12] to be usable in a marker assisted selection for high yield genotypes. These results suggested that DAPC is an appropriate method to analyze the organization of the genetic variation within and between mutant population in order to identify the best genotype as shown in our study, the individuals belonging to the same group shared several agromorphological characters. For instance, group 1 (Fig 6B) included white-seeded genotypes and long pod length unlike group 3 which showed short pod length.

## Conclusion

In this study, it was obviously noticed that gamma irradiation is a powerful tool to induce genetic variation in cowpea to broaden its genetic basis in order to overcome the bottleneck effect undergone by this crop during its domestication. This assertion has been supported by the various agromorphological variations affecting plant growth habit, flower color, pod length and weight, seed color, size and weight noticed among our mutant populations. These phenotypic variations were explained by the genetic variation revealed by 13 ISSR markers and the use of the datasets generated by these molecular markers enhanced our understanding on the mutant population structure. Overall, efficient exploitation of both agromorphological and molecular data led to the identification of promising high yielding new mutant genotypes which can be proposed for multi trial tests to assess their performance and stability in different agroecological conditions in Senegal and abroad but also their nutritional value. These mutant genotypes are valuable genetic resources usable for breeding programs but also for gene discoveries such as the ones involved in growth habit, pod length or seed size in cowpea.

## Acknowledgments

We thank the IAEA for funding this project. We thank also Thuloane Bernard TSEHLO and Dr Fatma SARSU respectively Programme Management Officer and Technical Officer for this project at the IAEA for their advices and support. I would also like to thank the anonymous reviewers for their valuable comments.

## Author Contributions

MD, FAB and DD designed the study. MD and SD performed DNA extraction and genotyping. MD and SD performed data analyses, OD run STRUCTURE and DAPC analyses. MD, SD, FAB and DD drafted the manuscript. All authors contributed to the final version.

## Supporting information

**S1 Table. Estimation of Pearson’s correlation between the agromorphological characters in the M6 of the mutant lines.**

**S2 Table. Estimation of Pearson’s correlation between the agromorphological characters in the M5 of the mutant lines.**

**S1 Fig. First Leave abbnormalities observed among the Melakh mutants.**

**A**: Two opposite leaves observed during the juvenile stage; **B**: Three leaves observed on the mutant

**C** and **D**: Leaves observed during adult stage on the mutant (tetraleaflet) and the control respectively

**S2 Fig. Variation of the flowering date after sowing (DAS) in the populations. A**: Melakh and its mutants; **B**: Yacine and its mutants

## Notes

### Competing Interest Statement

The authors have declared no competing interest.

