## Supplementary figures and images for "Development of new cowpea (*Vigna unguiculata*) mutant genotypes, analysis of their agromorphological variation, genetic diversity and Population structure"

### Supplementary Figure 1

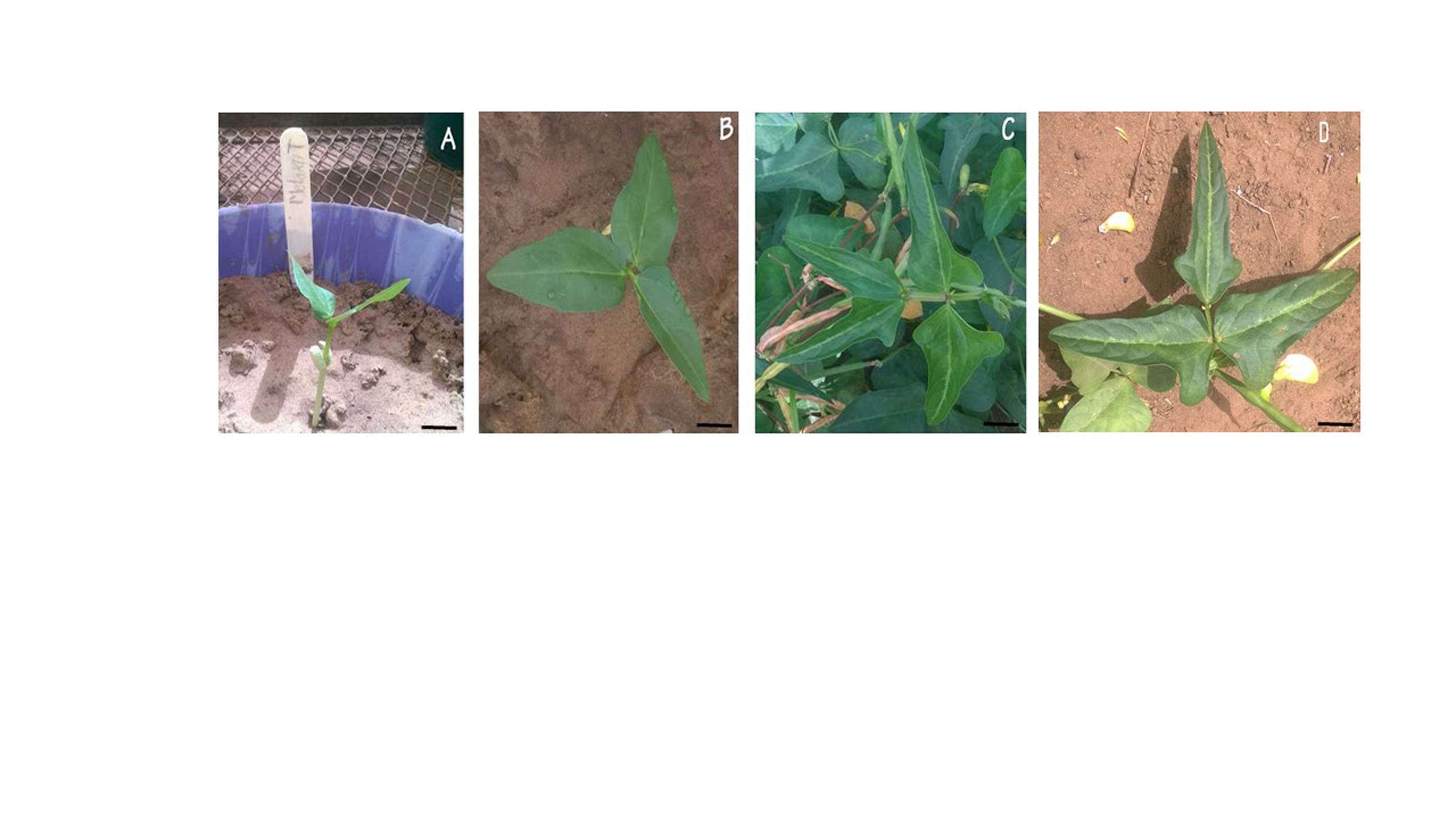

### Supplementary Figure 2

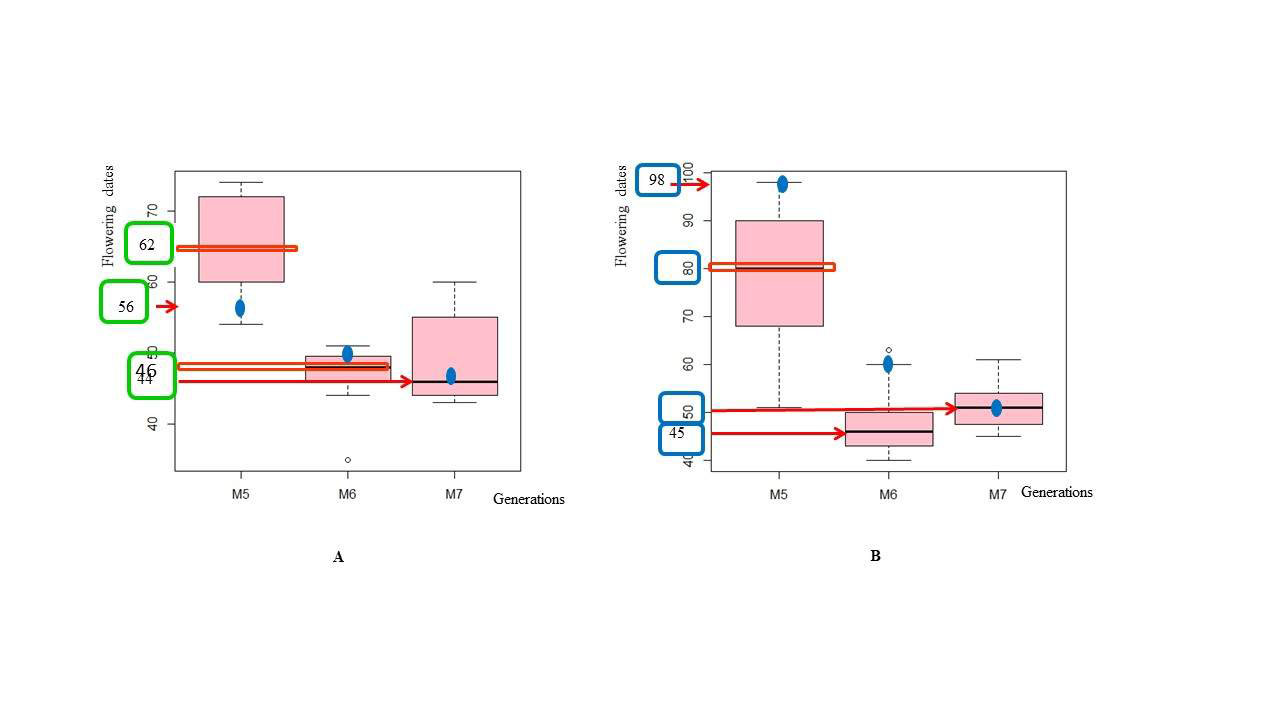
