## Supplementary Table 1 for "Development of new cowpea (*Vigna unguiculata*) mutant genotypes, analysis of their agromorphological variation, genetic diversity and Population structure"

**Supplementary Table S1: Estimation of Pearson’s correlation between the agromorphological characters in the M6 of the mutant lines**

|  | **PH** | **pig** | **GH** | **DF** | **FC** | **PdL** | **PdW** | **SL** | **SW** | **SWg** | **NSP** | **PdN** | **PdWg** | **SC** |
| --- | --- | --- | --- | --- | --- | --- | --- | --- | --- | --- | --- | --- | --- | --- |
| PH | 1 |  |  |  |  |  |  |  |  |  |  |  |  |  |
| pig | -0,01 | 1 |  |  |  |  |  |  |  |  |  |  |  |  |
| GH | 0,11 | -0,26 | 1 |  |  |  |  |  |  |  |  |  |  |  |
| DF | 0,28 | -0,15 | -0,16 | 1 |  |  |  |  |  |  |  |  |  |  |
| FC | -0,36* | 0,04 | 0,43** | -0,27 | 1 |  |  |  |  |  |  |  |  |  |
| PdL | 0,56*** | 0,15 | 0,13 | 0,28 | -0,33* | 1 |  |  |  |  |  |  |  |  |
| PdW | -0,11 | -0,19 | -0,04 | -0,02 | -0,05 | -0,24 | 1 |  |  |  |  |  |  |  |
| SL | 0,06 | -0,11 | 0,11 | -0,05 | 0,06 | -0,07 | 0,24 | 1 |  |  |  |  |  |  |
| SW | 0,10 | -0,25 | 0,49** | -0,12 | 0,19 | 0,03 | 0,16 | 0,31* | 1 |  |  |  |  |  |
| SWg | 0,15 | -0,17 | 0,04 | 0,07 | -0,19 | 0,29 | 0,17 | **0,60***** | 0,28 | 1 |  |  |  |  |
| NSP | 0,24 | 0,03 | 0,25 | 0,06 | -0,07 | **0,62***** | -0,13 | -0,10 | 0,16 | 0,29 | 1 |  |  |  |
| PdN | 0,32* | -0,06 | 0,68*** | -0,13 | 0,07 | 0,49** | -0,20 | -0,10 | 0,09 | 0,03 | 0,39* | 1 |  |  |
| PdWg | -0,19 | 0,01 | 0,06 | 0,03 | 0,07 | -0,06 | -0,22 | 0,21 | 0,10 | 0,03 | 0,11 | 0,03 | 1 |  |
| SC | 0,04 | **-0,59***** | 0,28 | 0,20 | 0,18 | -0,06 | -0,03 | 0,05 | 0,14 | 0,01 | 0,23 | -0,01 | 0,01 | 1 |

* significant at 5% level of probability; ** significant at 1% level of probability; *** significant at 0.1% level of probability

The number in bold indicate the agromorphological characters which are significantly correlated across generations
