## Supplementary Table 2 for "Development of new cowpea (*Vigna unguiculata*) mutant genotypes, analysis of their agromorphological variation, genetic diversity and Population structure"

**Supplementary Table S2: Estimation of Pearson’s correlation between the agromorphological characters in the M5 of the mutant lines**

|  | **PH** | **pig** | **GH** | **DF** | **FC** | **PdL** | **PdW** | **SL** | **SW** | **SWg** | **NSP** | **PdN** | **PdWg** | **SC** |
| --- | --- | --- | --- | --- | --- | --- | --- | --- | --- | --- | --- | --- | --- | --- |
| PH | 1 |  |  |  |  |  |  |  |  |  |  |  |  |  |
| pig | 0,19 | 1 |  |  |  |  |  |  |  |  |  |  |  |  |
| GH | 0,75*** | 0,14 | 1 |  |  |  |  |  |  |  |  |  |  |  |
| DF | -0,33 | 0,54*** | -0,50** | 1 |  |  |  |  |  |  |  |  |  |  |
| FC | -0,19 | 0,20 | -0,36* | 0,24 | 1 |  |  |  |  |  |  |  |  |  |
| PdL | 0,56*** | 0,18 | 0,54*** | -0,46** | -0,46** | 1 |  |  |  |  |  |  |  |  |
| PdW | 0,49** | 0,31 | 0,54*** | -0,44** | -0,29 | 0,76*** | 1 |  |  |  |  |  |  |  |
| SL | 0,54*** | 0,54*** | 0,49** | -0,50** | -0,13 | 0,64*** | 0,78*** | 1 |  |  |  |  |  |  |
| SW | 0,56*** | 0,31 | 0,42* | -0,26 | -0,17 | 0,66*** | 0,73*** | 0,75*** | 1 |  |  |  |  |  |
| SWg | 0,58** | 0,42* | 0,57*** | -0,50** | -0,18 | 0,76*** | 0,86*** | **0,86***** | 0,85*** | 1 |  |  |  |  |
| NSP | 0,40* | -0,08 | 0,39* | -0,03 | -0,09 | **0,42*** | 0,07 | 0,12 | 0,24 | 0,19 | 1 |  |  |  |
| PdN | -0,26 | -0,10 | -0,20 | -0,34* | -0,29 | 0,04 | 0,03 | -0,04 | -0,22 | -0,12 | -0,32 | 1 |  |  |
| PdWg | 0,53** | 0,06 | 0,51** | -0,22 | -0,33* | 0,48** | 0,40* | 0,37* | 0,48** | 0,52** | 0,47** | -0,26 | 1 |  |
| SC | -0,19 | **-0,50**** | -0,21 | 0,39* | 0,39* | -0,34* | -0,47** | -0,43* | -0,30 | -0,40* | 0,12 | -0,07 | -0,18 | 1 |

* significant at 5% level of probability; ** significant at 1% level of probability; *** significant at 0.1% level of probability

The number in bold indicate the agromorphological characters which are significantly correlated across generations
